# Integrating Human Genetics and Protective Genome Editing to Enable ADGRE2-Directed AML Therapy

**DOI:** 10.1101/2025.08.21.671614

**Authors:** Yonina Keschner, Julia Etchin, Mariana Silva, Hannah Mager, Hillary Hoyt, Juliana Xavier-Ferrucio, Nathan Manalo, Amanda Halfond, Huan Qiu, Ruijia Wang, Michelle I. Lin, Huanying Gary Ge, Julian Scherer, Tirtha Chakraborty, John R. Lydeard

## Abstract

Acute myeloid leukemia (AML) remains a major therapeutic challenge due to extensive disease heterogeneity and lack of cancer-specific antigens. ADGRE2 has emerged as a promising AML target with broad expression in AML patient blast and leukemic stem cell-enriched populations. However, comparable expression in healthy hematopoietic stem and progenitor cells (HSPCs) and myeloid lineages suggests a high susceptibility to on-target, off-tumor myelotoxicity with ADGRE2-targeted therapies. Guided by human genetics data identifying loss-of-function variants, we evaluated whether ADGRE2 is dispensable in hematopoietic stem cells as a protective approach for transplant-based shielding from ADGRE2-directed therapies. Using CRISPR-Cas9 and adenine base editors, we achieved high-efficiency ADGRE2 knockout (>94%) in HSPCs with corresponding protein loss without impairing cell viability, differentiation, and cytokine release *in vitro*, or long-term engraftment, multilineage differentiation, and persistence of gene editing in mouse xenografts. We also developed novel ADGRE2-specific chimeric antigen receptor (CAR) T cells that demonstrated potent cytotoxicity against AML cells, even at low antigen levels. Together, these findings establish ADGRE2 as a compelling AML target and provide a framework for hematopoietic stem cell transplant with protective gene editing to enable ADGRE2-directed immunotherapies while minimizing myelotoxicity.

## INTRODUCTION

Acute myeloid leukemia (AML) is an aggressive hematologic malignancy characterized by the clonal expansion of abnormal myeloid blasts that disrupt normal hematopoiesis (1,2). Although combination chemotherapy can induce complete remission in many patients, relapse is common, often occurring within 10 months (3). Efforts to improve outcomes have shifted toward targeted approaches, including small molecule inhibitors and antigen-specific immunotherapies, such as antibody drug conjugates, T cell engagers, and CAR T cells (4–6). While these strategies have shown promise (7), their clinical impact is limited by the substantial clonal heterogeneity of AML (8–10) and the scarcity of truly leukemia-specific antigens (11–13). Durable remission will likely require the eradication of both leukemic blasts and leukemia stem cells (LSCs) **—**the latter of which are thought to persist after therapy and drive disease relapse (8,9).

Allogeneic hematopoietic stem cell transplantation (allo-HCT) remains the most effective established, potentially curative therapy for many patients (3). Advances in conditioning regimens, supportive care, and the use of alternative donor sources have significantly expanded the applicability of allo-HCT (3,14,15). However, post-transplant relapse remains a major clinical challenge, especially in patients with adverse-risk disease or measurable residual disease at the time of transplant (3,16). Many antigen-directed therapies are limited by on-target, off-tumor myelotoxicity—due to antigen overlap between leukemic and normal hematopoietic cells. This has driven interest in “protective editing” strategies, in which donor-derived hematopoietic stem and progenitor cells (HSPCs) are engineered to remove the targeted antigen, allowing post-transplant immunotherapy to selectively eliminate residual leukemia while sparing healthy hematopoiesis (12,17,18).

We recently developed a comprehensive multimodal and longitudinal single-cell resource that quantifies expression intensity of 81 surface antigens and their co-expression patterns across individual leukemic cells, providing a detailed framework to guide immunotherapeutic strategies directed at AML heterogeneity (19). Among the top candidate targets—based on expression in AML and low-to-absent expression in healthy non-hematopoietic tissues—is the adhesion G protein-coupled receptor ADGRE2 (EMR2 or CD312). Previous studies have highlighted the expression of ADGRE2 in myeloid malignancies, and its therapeutic potential is currently being evaluated in a Phase I clinical trial employing a combinatorial logic-gated CAR T cell design (19–21).

Here, we evaluate ADGRE2 as a therapeutic target using mono-specific CAR T cell therapies, identified through high-throughput screening of candidate binders for specificity and affinity. Importantly, our surface expression analysis demonstrated comparable expression levels between AML populations (blasts and leukemic stem cell (LSC)-enriched populations) and several healthy hematopoietic populations (HSPCs and myeloid lineage cells), raising concerns for potential on-target, off-tumor effects in healthy hematopoietic cells upon ADGRE2 targeted-treatment. Because ADGRE2 is comparably expressed in leukemic and normal hematopoietic compartments, we evaluated whether protective editing could mitigate on-target myelotoxicity. Guided by human genetic loss-of function data, we used genome editing strategies to ablate *ADGRE2* in healthy donor HSPCs and evaluated functional consequences *in vitro* and *in vivo* to assess the feasibility of ADGRE2-protective editing for allo-HCT-based AML therapy.

## MATERIALS AND METHODS

### Primary samples

Patient derived AML bone marrow aspirates were collected with informed patient consent in accordance with the Declaration of Helsinki, and bone marrow mononuclear cells (BMMC) were isolated via standard Ficoll density gradient centrifugation; samples obtained cryopreserved from US BioLab (Germantown, MD, USA). Whole human bone marrow (BM) and non-mobilized Leukopak samples were obtained from healthy donors through StemExpress (Folsom, CA, USA). Cryopreserved CD34^+^ isolated cells from the peripheral blood of dual mobilized (G-CSF + Plerixafor) healthy donors were received from Charles River Laboratories (Wilmington, MA, USA) or StemExpress.

### Quantification of ADGRE2 Expression

ADGRE2 surface expression was quantified using a Phycoerythrin (PE)-conjugated anti-ADGRE2 antibody (clone REA301 or 2A1; Miltenyi Biotec Inc., Charlestown, MA, USA) and QuantiBRITE^TM^ beads (BD Biosciences, Franklin Lakes, NJ, USA) via flow cytometry following manufacturer instructions. Interpolation of geometric mean fluorescence intensity values was completed on GraphPad Prism (for MacOS).

### CAR generation and cytotoxicity

Ten novel ADGRE2-directed binders were identified by phage display panning using Navi-His ADGRE2 against human single-chain variable fragment (scFv) and heavy chain variable region (VH) libraries, as previously described in WO2024/238565 (22). Specific ADGRE2-binding was evaluated through multi-point flow cytometric analysis using MOLM-13 wild-type (WT) and HEK-293 cells, as well as by Enzyme-Linked Immunosorbent Assay (ELISA) and Blitz analyses. Lack of cross reactivity with CD97 was confirmed by ELISA. Selected binders were used to generate six, second generation ADGRE2-directed CAR constructs with a 4-1BB co-stimulatory domain. CAR constructs were delivered into primary T cells via lentiviral transduction. Effectors (E) and CFSE-labeled Targets (T) were co-cultured in triplicate at indicated E:T ratios for 24 or 48 hours. Cells were harvested for flow cytometry and supernatant was collected for cytokine analysis.

### Construction of expression vectors

For CAR binding studies, expression DNA constructs were cloned into the pcDNA3.1(+)-P2A-eGFP backbone vector (GenScript, Piscataway, NJ, USA). For studies of naturally occurring ADGRE2 variants, constructs were cloned into the pcDNA3.1(+)-IRES-eGFP backbone vector (GenScript) based on the ADGRE2 sequence (CCDS32935.1); an N-terminal HA epitope tag (YPYDVPDYA) was inserted directly downstream of the native signal peptide.

### Engineered Antigen Diverse Clones

MOLM-13 ADGRE2-knockout (KO) cells (CRISPR-Cas9) were engineered to stably express a range of variable but distinct ADGRE2 surface antigen molecules. Lentiviral vectors were constructed using the ADGRE2 coding sequence (CCDS32935.1) into the pGenLenti backbone (GenScript). DNA plasmids were transfected into HEK293F cells (Thermo Fisher Scientific, Waltham, MA, USA), followed by lentiviral transduction in MOLM-13 ADGRE2-KO cells and single-cell sorting.

### Gene Editing in CD34^+^ HSPCs

*ADGRE2* gene disruption was achieved in human CD34^+^ HSPCs with either CRISPR-Cas9 RNP, adenine base editing (ABE), or cytosine BE (CBE) to induce protein KO or disruption via electroporation (EP). Further details on guides used for KO can be found in the associated patent WO2023086422 (23).

### *In-vitro* myeloid differentiation and functional assessment of HSPCs

Forty-eight hours post-EP, ADGRE2-KO and non-edited, control CD34^+^ HSPCs were transferred into monocytic-differentiating culture conditions (STEMCELL Technologies, Cambridge, MA, USA) and evaluated for viability, gene editing, phenotypic surface expression analysis, and cytokine release as previously described (12). A complete list of antibodies can be found in Supplemental Table 6.

### Analysis of HSPC sub-populations

CD34^+^ HSPCs were electroporated, and after 48 hours were stained with flow cytometry techniques and bulk sorted into defined subpopulations based on established surface marker profiles (24,25).

### In vivo pharmacology

Xenotransplant NOD-scid IL2Rg null (NSG) mice were injected with ADGRE2-KO or non-edited CD34^+^ HSPCs. Bone marrow and peripheral blood were analyzed by flow cytometry at the endpoint. Editing efficiency was measured pre-infusion and persistence of editing was assessed in engrafted mice bone marrow by next-generation sequencing (rhAmpSeq, Integrated DNA Technologies, Coralville, IA, USA).

See Supplemental Data for a detailed Materials and Methods section.

## RESULTS

### ADGRE2 is expressed on primary AML blasts and leukemic stem cells

Previous studies have highlighted ADGRE2 as a promising AML target due to its broad expression across primary AML samples (20,26,27). To confirm and extend these observations, we performed flow cytometric analysis on primary AML BMMCs (Figure 1A). To assess therapeutic relevance of ADGRE2 expression, surface levels of ADGRE2^+^ cells were quantified in AML blast and leukemic stem cell (LSC)-enriched compartments (28,29) using a standard curve created by QuantiBRITE beads. ADGRE2 was detected in all patient samples, with a median 94.3% expression in AML blasts (range: 18.3%-99.7%) and 98.0% in the LSC compartment (46.9%-100.0%). Among ADGRE2-expressing populations, the median antigen intensity was 1,059 antigens per cell (APC) in AML blasts (range: 391-3134) and 969 in LSCs (294–3730). These findings confirm widespread ADGRE2 surface expression across AML patient samples and support its potential as an immunotherapy target (30–32).

**Figure 1.**
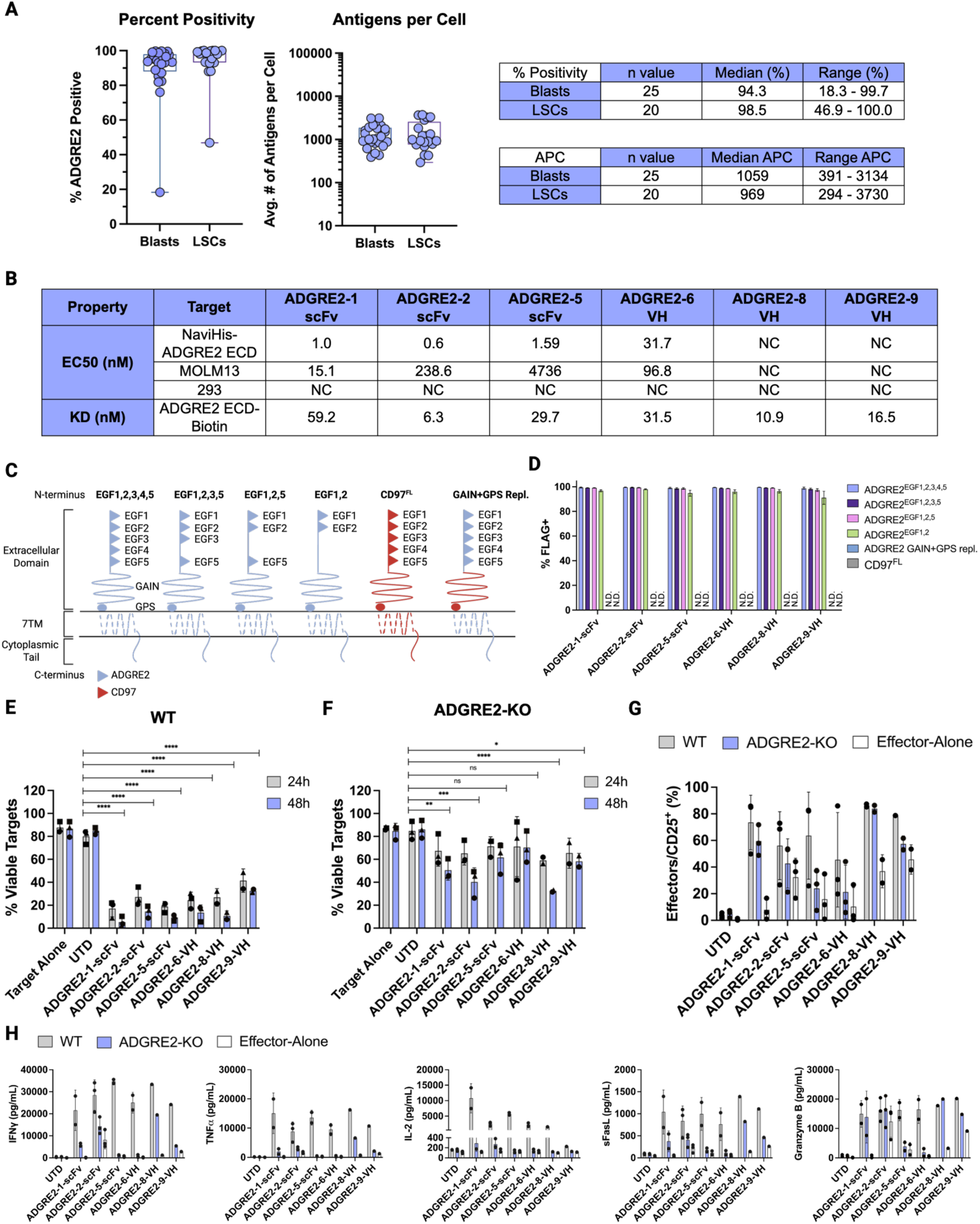
Characterization of ADGRE2 Expression in AML and Functional Evaluation of ADGRE2-Directed CAR Ts. **(A)** Flow cytometric analysis of ADGRE2 surface expression and number of antigens per cell (APC) on primary AML bone marrow samples. Data shown for AML blasts (CD45^dim^SSC^lo^; n=25) and leukemic stem cell (LSC)-enriched compartments (CD34^+^CD38^-^; n=20). Each data point represents an individual patient. Quantification of ADGRE2 expression was performed using a standard curve generated by QuantiBRITE beads and interpolation of geometric mean fluorescence intensity values of positively expressing cells. Data represented as box-and-whisker plots indicating the 25^th^, 50^th^ (median), and 75^th^ percentiles; whiskers denote minimum and maximum values. **(B)** EMR2-directed binders were identified by phage display panning against single-chain variable fragment (scFv) and heavy chain variable region (VH) libraries. Binder kinetic and affinity characterization (EC50 and KD) were performed by multipoint flow cytometric, ELISA and Blitz analyses. **(C)** Schematic representation of ADGRE2 full-length (FL) and splice isoforms, FL CD97, and a chimeric construct in which the native ADGRE2 GAIN and GPS domains were replaced with the corresponding regions from CD97. Illustration created using BioRender. **(D)** Binding specificity of ADGRE2-directed CAR binders to ADGRE2 isoforms, FL CD97, and the chimeric ADGRE2-CD97 construct (GAIN and GPS replacement) expressed on the surface of HEK293T cells following transient transfection. CAR constructs were tagged with a C-terminal FLAG peptide; binding was assessed by flow cytometry and anti-FLAG secondary detection. **(E-F)** *In vitro* cytotoxicity of ADGRE2-directed CAR Ts (ADGRE2-1-scFv, ADGRE2-2-scFv, ADGRE2-5-scFv, ADGRE2-6-VH, ADGRE2-8-VH, or ADGRE2-9-VH) or untransduced T cells (UTD) against MOLM-13 wild-type (WT) and ADGRE2-knockout (KO) targets at an effector-to-target (E:T) ratio of 1:1. Cytotoxicity assessed at 24h and 48h by flow cytometry; viable targets defined as Annexin-V^-^/LIVE-DEAD^-^. Each data point represents an individual T cell donor; data shown as mean ± SD from 2-3 donors. Two-way ANOVA with Šidák multiple comparisons test was performed to compare viability of targets in co-culture with CAR T cells to UTD, averaging both timepoints; *p < 0.05, **<0.01, ***p < 0.001, ****p<0.0001. **(G)** T cell activation measured by CD25 surface expression after 48h co-culture. UTD and effector-alone conditions served as controls. Data shown as mean ± SD from 2-3 donors; each data point represents a donor. **(H)** Cytokine secretion profile of ADGRE2-directed CAR Ts and UTD controls following 48 h co-culture with MOLM-13 WT or ADGRE2-KO targets at a 1:1 E:T ratio. Supernatants were analyzed for IFNγ, TNFα, IL-2, sFasL, and Granzyme B. Effector-alone controls included. Data shown as mean ± SD from 1-3 donors, each dot represents a donor. Abbreviations: scFv, single-chain variable fragment; VH, heavy chain variable domain; GAIN, G-protein-coupled receptor (GPCR) autoproteolysis-inducing domain; GPS, GPCR proteolytic site; 7TM, 7-pass transmembrane domain; SD, standard deviation; ns, not significant; IFNγ, interferon gamma; TNFα, tumor necrosis factor alpha; IL-2, interleukin-2; sFasL, soluble Fas ligand; N.D., not detected.

### ADGRE2-specific binders recognize a native GAIN/GPS-dependent epitope

To evaluate the targetability of ADGRE2, we developed novel ADGRE2-directed CAR T binders using phage display panning against human scFv and VH libraries. Binder characterization was performed by flow cytometry, ELISA-based EC50, and Blitz-based KD measurements. Ten binders were initially screened; three ScFv and three VHs were selected based on specific binding to recombinant ADGRE2 protein and ADGRE2-expressing MOLM-13 cells (Figure 1B), with no cross-reactivity to CD97 (ADGRE5; a closely related adhesion GPCR with high sequence homology to ADGRE2 (33); data not shown). All six binders demonstrated robust binding (>85%) across common ADGRE2 isoforms (34–36), as measured using anti-FLAG secondary detection (Figures 1C,1D). Lack of binding to CD97 was confirmed, supporting binder specificity. To further characterize binder specificity, we mapped the general epitope region recognized by the six ADGRE2-targeting constructs. A chimeric ADGRE2 molecule was engineered by replacing the native GPCR autoproteolysis inducing (GAIN) domain and GPCR proteolytic site (GPS) with the corresponding regions from CD97. The domain swap abrogated binding for all six constructs, indicating that recognition is dependent on the native ADGRE2 GAIN/GPS region.

### ADGRE2-directed CAR T cells exhibit potent, selective AML cytotoxicity

To assess the cytotoxic potential and specificity of ADGRE2-directed CAR T cells against AML, we performed *in vitro* co-culture assays using MOLM-13 targets. All ADGRE2-directed CAR T effectors induced significant target cell killing against wild-type (WT) MOLM-13 within 24 hours, with viability declining to <33% by 48 hours (Figure 1E). Untransduced T cells (UTD) had no effect on viability, confirming CAR-dependent killing. However, off-target assessment with ADGRE2-KO MOLM-13 revealed varying degrees of non-specific activity (Figure 1F)— moderate reduction in viability with constructs ADGRE2-1-scFv, ADGRE2-2-scFv, and ADGRE2-8-VH (36%, 46%, and 54% compared to UTD at 48 hours, respectively), whereas ADGRE2-5-scFv, ADGRE2-6-VH, and ADGRE2-9-VH showed minimal off-target effects (25%, 16%, and 28% reduction, respectively). Importantly, ADGRE2-5-scFv and ADGRE2-6-VH also demonstrated minimal background T cell activation or cytokine release when exposed to KO targets or when cultured alone (Figures 1G, 1H). Based on its favorable profile of specific cell killing, T cell activation and cytokine release, ADGRE2-5-scFv was selected as the lead CAR T construct for further characterization.

### ADGRE2-5-scFv CAR T demonstrates sensitivity to low antigen expression

Broad ADGRE2 surface expression on AML populations signals the need to define the minimum antigen threshold for effective ADGRE2-5-scFv activity. To investigate this, we generated two clonal MOLM-13 cell lines stably expressing reduced ADGRE2 surface levels compared to WT cells (Figure 2A). These lines were co-cultured with ADGRE2-5-scFv CAR T cells, generated from three independent donors (Figures 2B, 2C; Supplemental Figure 1A); CAR T and UTD effector cells were characterized for CD4 and CD8 expression (Supplemental Figures 1B, 1C).

**Figure 2.**
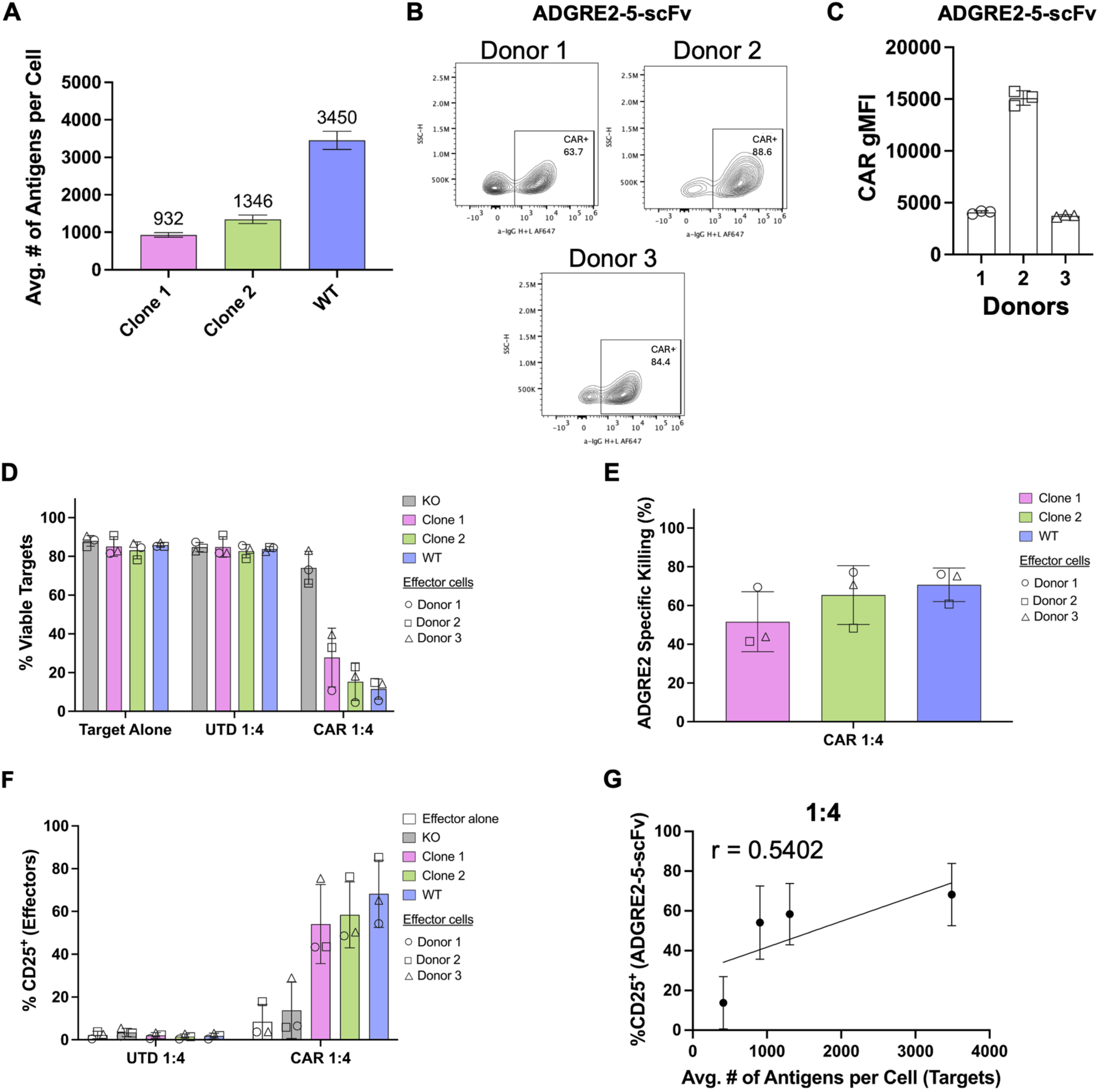
ADGRE2-5-scFv CAR T cells mediate cytotoxicity against target cells expressing low levels of ADGRE2. **(A)** Flow cytometric analysis of ADGRE2 surface expression on MOLM-13 wild-type (WT) and lentivirus engineered clones. ADGRE2 surface quantification was completed using QuantiBRITE bead interpolation. Data shown as mean ± SD from n=3 independent measurements. **(B)** Representative flow cytometry plots of ADGRE2-5-scFv CAR surface expression (anti-human IgG H+L) from n=3 T cell donors. **(C)** Geometric mean fluorescence intensity (gMFI) of ADGRE2-5-scFv CAR expression in transduced T cells from panel (B). Data shown as mean ± SD; each data point represents a technical replicate (n=3). **(D)** *In vitro* cytotoxicity assessment of untransduced (UTD) or ADGRE2-5-scFv (CAR) T cells against MOLM-13 ADGRE2-KO, Clone 1, Clone 2, or WT targets at a 1:4 effector-to-target (E:T) ratio. Viability was assessed after 48 h by flow cytometry (Annexin-V^-^/LIVE-DEAD^-^). Target-alone cultures served as controls. Data represented as mean ± SD from n=3 T cell donors. **(E)** ADGRE2-specific killing of MOLM-13 clones and WT at a 1:4 E:T ratio, calculated as (*Viability_untreated_* – *Viability_treated_*)/*Viability_untreated_ x* 100 followed by subtracting killing observed in KO targets for background correction. Data represented as mean ± SD from n=3 T cell donors. **(F)** Effector activation measured by CD25 surface expression after 48 h co-culture with indicated MOLM-13 targets; flow cytometry analysis. Effector-alone condition included as a control. Data shown as mean ± SD from n=3 donors. **(G)** Correlation between ADGRE2-5-scFv T cell activation (CD25^+^) and ADGRE2 surface intensity (antigens per cell, APC) on MOLM-13 targets at a 1:4 effector-to-target (E:T) ratio. Data shown as mean ± SD. Pearson correlation efficient (r) and linear regression analysis were generated using GraphPad Prism. Abbreviations: SD, standard deviation; KO, knockout.

After 48-hours, ADGRE2-5-scFv CAR Ts demonstrated robust, antigen-dependent cytotoxicity against MOLM-13 cells expressing as few as ∼932 ADGRE2 surface molecules. At an effector-to-target (E:T) ratio of 1:4, <40% of viable targets remained (Figure 2D), whereas UTD controls showed minimal cytotoxicity. Non-specific killing of ADGRE2-KO target cells was <11% at an E:T of 1:4, resulting in antigen-specific cytotoxicity of 52-71% for all ADGRE2 expressing MOLM-13 targets (Figure 2E). Effector activation of ADGRE2-5-scFv CAR T cells, measured by CD25 expression, exceeded an average of 40% in co-cultures with all ADGRE2-expressing targets, but remained low in KO co-cultures and with UTD effectors (Figure 2F). Furthermore, CD25 expression is positively correlated with ADGRE2 levels on target cells, indicating activation is antigen-dependent (Figure 2G).

At a higher E:T of 1:1, ADGRE2-5-scFv CAR Ts exhibited enhanced cytotoxic activity, reducing viability of all MOLM-13 lines to below 30% (Supplemental Figure 1D). This increased activity was accompanied by higher levels of non-specific anti-leukemia effects across targets. After background correction for KO-target killing, antigen-specific cytotoxicity ranged from 20-38% (Supplemental Figure 1E). Effector activation at this higher E:T mirrored trends seen at 1:4, but showed modestly increased CD25 expression in KO co-cultures versus effectors alone, suggesting potential non-specific activation at elevated effector levels (Supplemental Figures 1F, 1G). In summary, ADGRE2-5-scFv CAR Ts exhibit potent, antigen-specific cytotoxicity and activation at low antigen densities (<1000 surface molecules), suggesting promise for targeting AML blasts and leukemic stem cells.

### Activated T cells upregulate ADGRE2 surface expression

Previous studies offer conflicting evidence on ADGRE2 expression in activated lymphocytes (24,35,36). Given the fratricide risk in CAR T therapy (37) upon antigen-specific activation, we assessed ADGRE2 surface expression on activated T cells under process-relevant settings (Supplemental Figures 2A). Following a 48-hour co-culture with ADGRE2-expressing MOLM-13 targets, up to 36.1% of ADGRE2-5-scFv effectors showed increased ADGRE2 surface expression, positively correlating with target antigen intensity (Supplemental Figures 2B, 2C). Additionally, CD3/CD28-activated pan-T cells (>85% CD25/CD69 expression) exhibited a ∼10-15% increase of ADGRE2 surface expression in 2 out of 3 donors versus non-activated controls (Supplemental Figures 2D, 2E). These results suggest that activation can induce ADGRE2 upregulation in a subset of T cells. However, further studies are needed to clarify its function and assess fratricide risk for clinical mitigation.

### ADGRE2 expression is primarily restricted to healthy HSPCs and myeloid lineages

ADGRE2 expression is known to be largely restricted to hematopoietic cells, with evidence suggesting preferential expression in differentiated myeloid populations (38,39). To evaluate the potential for on-target, off tumor toxicity upon targeted treatment, we profiled ADGRE2 expression in healthy hematopoietic populations. In bone marrow-derived HSPC populations, ADGRE2 expression was broadly expressed across hematopoietic stem and progenitor populations (Figure 3A). Median ADGRE2 surface intensity across these populations ranged from 597 to 1,267 antigens per cell (APC), with the highest intensity observed in CLPs. In peripheral blood, mature myeloid populations exhibited high ADGRE2 expression with antigen intensity varying by lineage. ADGRE2 was expressed on 59.5% of neutrophils and on over 80% of the examined myeloid lineage cell types. Neutrophils exhibited the highest median APC (5,174), followed by cDCs (4,704), and basophils (3,285). By comparison, mature lymphoid populations showed minimal ADGRE2 expression (<15%) and low APC values (a distinction from the elevated expression observed in activated T cells; Supplementary Figures 2D, 2E).

**Figure 3.**
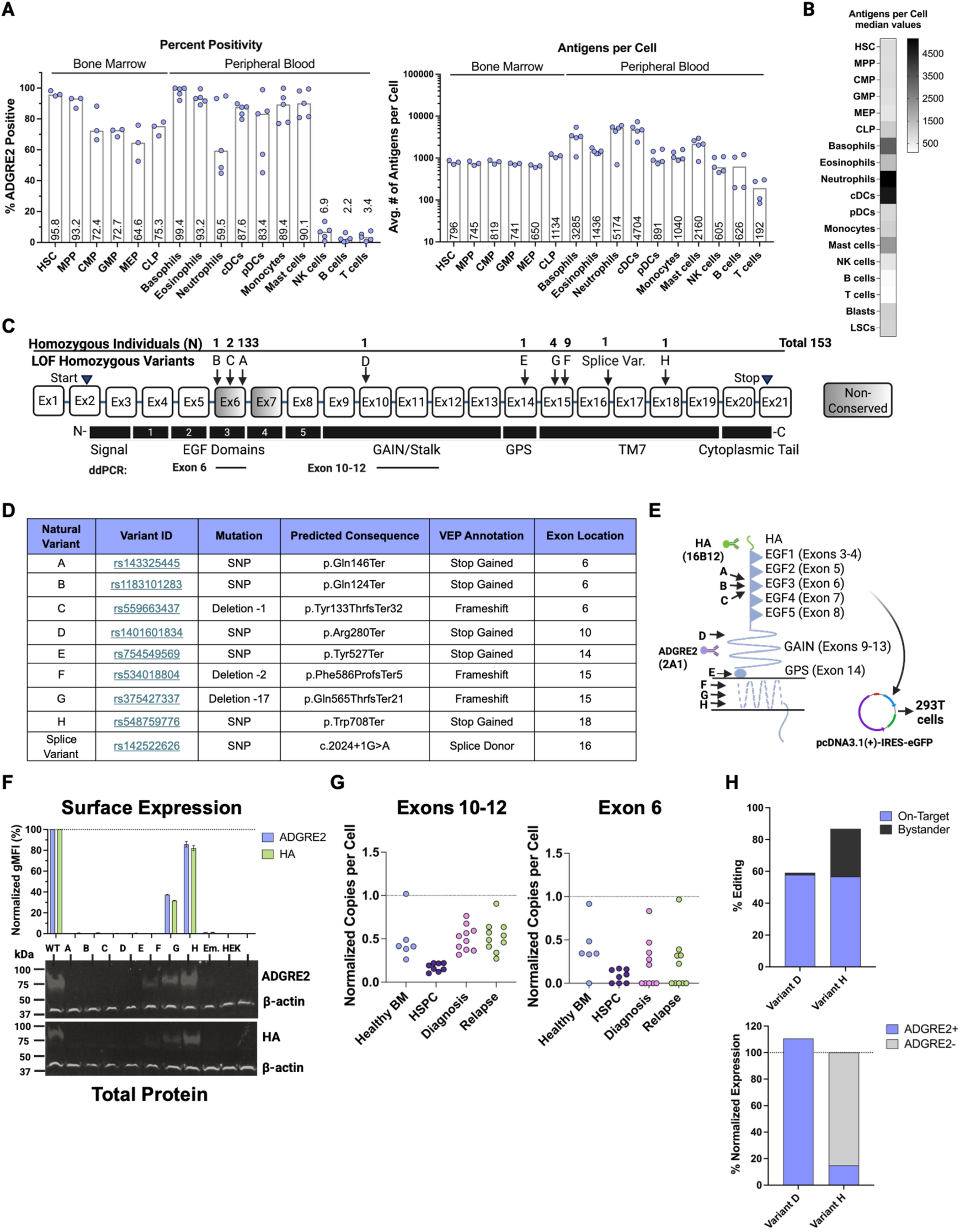
ADGRE2 Expression in Healthy Hematopoietic Compartments and Functional Validation of Human Loss-of-Function Variants. **(A)** Flow cytometric analysis of ADGRE2 percent surface expression and antigens per cell (APC) on healthy bone marrow (n=3) and peripheral blood (n=5) hematopoietic cell subsets. Quantification of ADGRE2 expression was performed by interpolation of PE geometric mean fluorescence intensity (gMFI) values using a QuantiBRITE bead-generated standard curve. Data represented as median values; each point represents an individual donor sample. One donor showed undetectable ADGRE2 expression on lymphocytes by QuantiBRITE interpolation. **(B)** Heatmap summarizing median ADGRE2 APC across healthy hematopoietic and AML disease populations. **(C)** Schematic of *ADGRE2* gene structure (not drawn to scale) with indicated start/stop codons (triangles) and associated coding regions. Positions of reported homozygous predicted loss-of-function (pLOF) variants and the number of affected individuals are indicated (arrows). Exonic regions analyzed by ddPCR are also shown. **(D)** Summary table of ADGRE2 homozygous pLOF natural variants identified in gnomAD v4.1.0. **(E)** Schematic of full-length ADGRE2 protein with indicated locations of pLOF variants (A-H; arrows). Wild-type (WT) and variant coding sequences were cloned into pcDNA3.1(+)-IRES-eGFP expression plasmids, with an N-terminal HA tag inserted in each construct. General binding regions for the anti-ADGRE2 (clone 2A1) and anti-HA antibody (clone 16B12) used in flow cytometry and Western blotting are shown. **(F)** Surface expression (geometric mean fluorescence intensity, gMFI) of ADGRE2 and HA in HEK293T cells transfected with WT and variant (A-H) ADGRE2 plasmids; measured by flow cytometry 24 h post-transfection and normalized to WT. Empty vector (em.) and non-transfected cells served as controls. Data represented as mean ± SD; n=2 biological replicates. Representative Western blot of total ADGRE2 and HA protein expression; β-actin served as a loading control. **(G)** Transcript expression of ADGRE2 exons 10-12 and exon 6 in healthy bone marrow (BM; n=6), CD34^+^ hematopoietic stem and progenitor cells (HSPC) day 2 post-thaw (n=9), and AML bone marrow samples at diagnosis (n=10) and relapse (n=10); measured by digital droplet PCR (ddPCR) and normalized to *GUSB* (glucuronidase beta) for each sample. **(H)** Base editing efficiency of Variants D and H in CD34^+^ HSPCs using BE4-PpAPOBEC1 or BE4-PpAPOBEC-SpG guide-RNAs targeting respective variant regions. Editing was assessed on day 5 post-electroporation by next-generation-sequencing (rhAmpSeq). “On-Target” represents reads with the intended C-to-T edit and no bystander conversions; “Bystander” includes reads with off-target C-to-T conversions alone or in combination with the intended on-target edit. ADGRE2 surface expression was assessed in parallel by flow cytometry and normalized to non-edited control HSPC cells; n=1 biological replicate. Abbreviations: HSC, hematopoietic stem cells; MPP, multipotent progenitor; CMP, common myeloid progenitor; GMP, granulocyte-monocyte progenitor; MEP, megakaryocyte-erythroid progenitor; CLP, common lymphoid progenitor; cDCs, classical dendritic cell; pDCs, plasmacytoid dendritic cells; NK, natural killer cell; HSPC, hematopoietic stem and progenitor cells; ddPCR, digital droplet PCR; SD, standard deviation.

These data confirm that ADGRE2 is expressed in HSPCs and myeloid lineages (33,40,41). Importantly, expression levels in healthy myeloid and progenitor compartments mirror those in AML blasts and LSCs (Figure 3B), highlighting the high risk for on-target myelotoxicity from ADGRE2-directed therapies. Protective strategies will likely be required to shield the hematopoietic compartment from these therapies.

### Human genetic and functional evidence suggest ADGRE2 is dispensable

To unlock the full potential of ADGRE2-targeted strategies, emerging protective editing approaches—such as those described to enable CD33-directed therapies (12)—can be leveraged to ablate ADGRE2 expression specifically in healthy hematopoietic cells, thereby shielding normal hematopoiesis while maintaining potent immunotherapeutic activity against AML. To assess the feasibility of this approach and mitigate potential myelotoxicities from ADGRE2-directed therapies, we first examined ADGRE2 population genetics data. Analysis of the human genetics database gnomAD (v4.1.0; (42)) revealed that ADGRE2 has a predicted loss-of-function intolerance (pLI) score of 0 and a loss-of-function observed/expected upper bound fraction (LOEUF) score of 1.03, indicating minimal evolutionary constraint. Notably, 153 individuals carry nine distinct homozygous predicted loss-of-function (pLOF) variants (Figures 3C, 3D).

To investigate the functional impact of these variants, we expressed each pLOF variant in HEK293T cells with N-terminal HA tags to enable orthogonal detection of ADGRE2, independent of antibody epitope recognition (Figure 3E). WT ADGRE2, empty vector (Em.), and non-transfected cells served as controls. Transfection efficiency, measured by eGFP expression, ranged from 34-48% across constructs (data not shown). Flow cytometry and Western blot analysis revealed that six variants (A-D and H) led to complete or near-complete loss of surface and total protein expression (Figure 3F); validating potential LOF alleles as bona fide protein KOs.

Variants A-C, located within exon 6, represent the majority of individuals with homozygous pLOF variants and showed absent expression; however, there is evidence of heterogeneous inclusion of exon 6 (EGF3) from alternative splicing in naturally occurring isoforms (34–36,43). Thus, while pLOF variants in exon 6 are common in gnomAD, alternative splicing likely reduces their functional impact, underscoring the need for functional validation. We therefore performed transcript-level analysis using ddPCR on healthy BM, CD34^+^ HSPC donor cells, and AML BMMC samples at both diagnosis and relapse (primer-probe locations mapped in Figure 3C). While total ADGRE2 transcript (assessed by expression of exons 10-12) was broadly expressed across all samples, exon 6 levels varied widely, with some samples displaying no detectable levels (Figure 3G). This suggests that Variants A-C may not represent true functional KOs as these exons are variably spliced out of the full protein.

In contrast, Variants D and H affected conserved regions of ADGRE2 (exons 14 and 18, respectively) and demonstrated robust loss of ADGRE2 surface and total protein, nominating them for further study. To confirm their functional impact in a physiologic context, we introduced the corresponding C-to-T single nucleotide polymorphisms (SNPs) for Variants D and H in CD34^+^ HSPCs using BE4-PpAPOBEC base editors. In a single CD34^+^ HSPC donor, next-generation sequencing revealed efficient (∼60%) on-target editing for both variants tested; Variant H also exhibited ∼30% bystander edits (Figure 3H). Surface protein loss was confirmed for Variant H (p.Trp708Ter), but not for Variant D (p.Arg280Ter) in CD34^+^ HSPCs. These findings highlight the importance of functional validation for pLOF variants, and the loss of protein observed following Variant H editing suggests that ADGRE2 is likely biologically dispensable for human hematopoiesis.

### ADGRE2 deletion does not impair monocytic differentiation of human HSPCs

To evaluate the impact of *ADGRE2* deletion on myelopoiesis, CRISPR-Cas9-mediated KO HSPCs were cultured under monocytic-differentiating conditions for 14 days alongside non-edited controls. Editing efficiency by ICE analysis (44) averaged 76.5% at day 2 post-electroporation and increased to >94% by day 6, resulting in sustained ADGRE2 surface loss through endpoint (Figure 4A). Viability and proliferation remained unaffected (Figure 4B). Differentiated cells from ADGRE2-KO HSPCs showed comparable expression of key myeloid markers (Figure 4C) and similar cytokine release (Figure 4D) relative to controls, indicating intact monocytic differentiation and function. However, analysis of insertion/deletion (InDel) patterns revealed a dominant -4 bp deletion that is suggestive of microhomology-mediated end joining (MMEJ) repair (data not shown; (45,46)). Given the limited persistence of MMEJ edits in long-term hematopoietic stem cells (LT-HSCs; (47)), we next evaluated base editor strategies to ablate *ADGRE2* in HSPCs.

**Figure 4.**
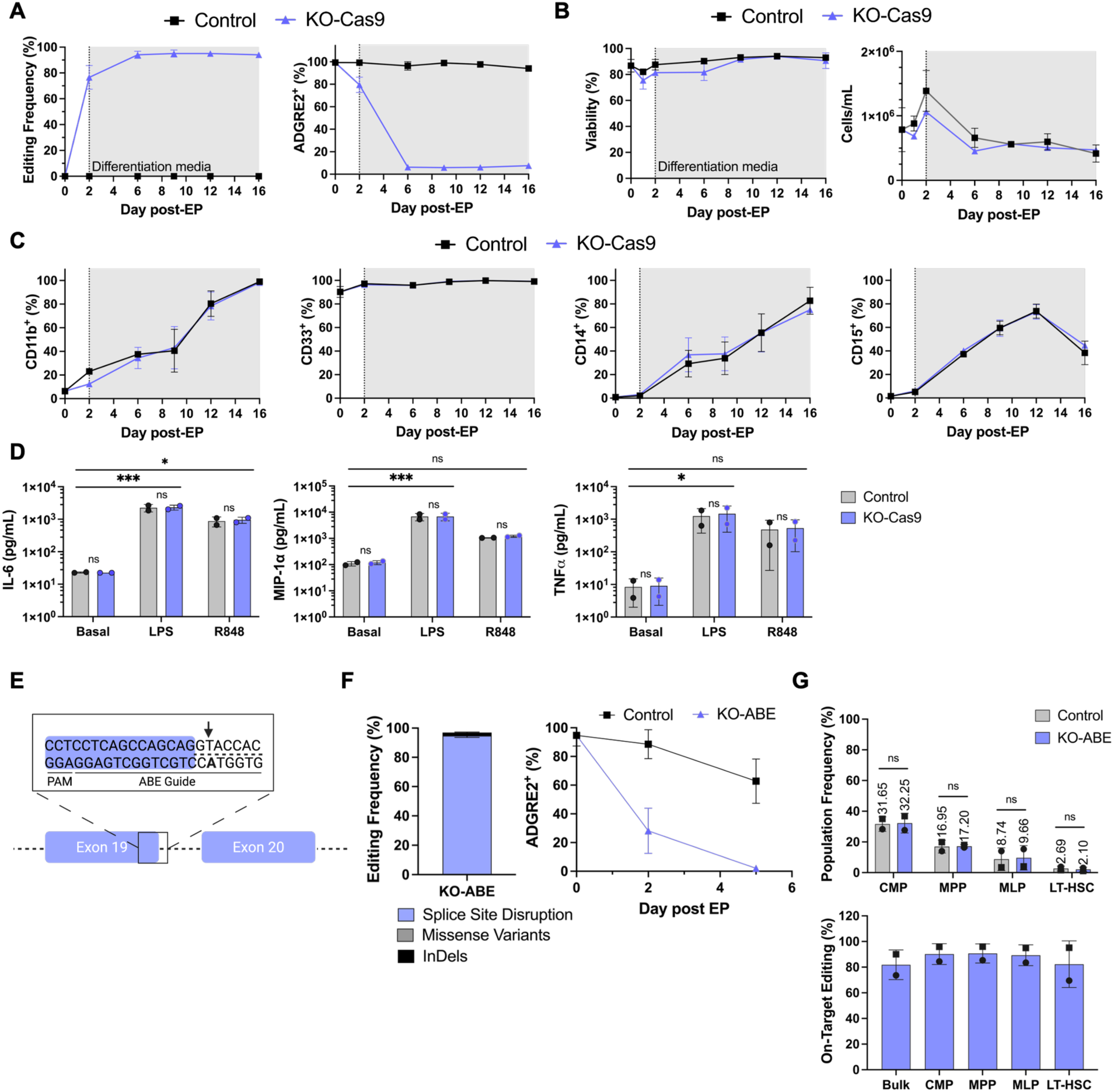
ADGRE2 deletion in human HSPCs does not impact myeloid differentiation or function *in vitro*. **(A)** CD34^+^ hematopoietic stem and progenitor cells (HSPCs) were electroporated with ADGRE2-targeting CRISPR-Cas9-ribonucleoprotein (RNP) complex (KO-Cas9) or a non-editing control RNP (Control). Two days post-electroporation (EP), cells were cultured for 14 days under monocytic differentiation conditions (indicated by dashed line and shaded grey). Editing frequency was assessed by ICE analysis and ADGRE2 surface expression was measured by flow cytometry. Data shown as mean ± SD; n=2 donors. **(B)** Cell viability and concentration throughout the 16-day *in vitro* culture. Quantified by cell counter; data shown as mean ± SD; n=2 donors. **(C)** Phenotypic analysis of differentiating cells by flow cytometry using CD11b (pan-myeloid marker), CD33 (myeloid marker), CD14 (monocytic lineage marker), and CD15 (granulocytic lineage marker). Data shown as mean ± SD; n=2 donors. **(D)** Inflammatory cytokine secretion (IL-6, MIP-1α, TNFα) evaluated in monocytic differentiated cells, collected from supernatant at study end. Data shown as mean ± SD; each dot represents a donor. Statistical analysis was performed using two-way ANOVA comparing basal condition versus LPS stimulation, or basal condition versus R848 (TLR7/8 agonist) stimulation; *p < 0.05, ***p < 0.001. Two-way ANOVA with Šidák multiple comparisons test was performed to compare KO-Cas9 to Control for each stimulation condition. **(E)** Schematic representation of the lead ABE8.20-m guide-RNA (gRNA) targeting the splice donor site at the end of exon 19 of *ADGRE2*. On-target A-to-G editing site is bolded and indicated by an arrow. **(F)** CD34^+^ HSPC were electroporated with ABE8.20-m mRNA and either an *ADGRE2*-targeting gRNA (KO-ABE) or non-targeting control (Control) and cultured for 5 days. Editing outcomes—splice-site disruption (intended A-to-G edit), missense conversions, and insertions/deletions (InDels)—were quantified by next-generation sequencing (rhAmpSeq). ADGRE2 surface protein expression was measured by flow cytometry. Data represented as mean ± SD; n=4 donors. **(G)** Frequency of HSPC subpopulations 48 h post-electroporation (KO-ABE and Control), measured by flow cytometry. Subsets included CMPs, MPPs, MLPs, and LT-HSCs, expressed as percentage of total live cells. Cells were bulk sorted and editing frequency of splice-site disruption was assessed by rhAmpSeq. Data shown as mean ± SD; n=2 donors. Statistical analysis by ANOVA with Šidák multiple comparisons test. Abbreviations: KO, knockout; SD, standard deviation; IL-6, interleukin-6; MIP-1α, macrophage inflammatory protein-1 alpha; TNFα, tumor necrosis factor alpha; LPS, lipopolysaccharide; TLR, toll-like receptor; CMP, common myeloid progenitors; MPP, multipotent progenitors; MLP, multi-lymphoid progenitors; LT-HSCs, long-term hematopoietic stem cells.

### ADGRE2 KO did not affect HSPC subpopulation distribution

Adenine base editors (ABEs) enable precise A-to-G edits without introducing double-stranded breaks, offering an alternative strategy to achieve durable gene disruption in hematopoietic stem cells (48). Guide RNA (gRNA) screening in CD34^+^ HSPCs identified a lead gRNA targeting the exon 19 splice site that disrupts ADGRE2 surface expression, achieving an average on-target editing of 94.8% (Figure 4E, 4F). Editing was accurate, with minimal bystander edits or InDels. To evaluate the biological impact of ADGRE2 editing outcomes across HSPC subpopulations, including LT-HSC, CMPs, MPPs, and multi-lymphoid progenitors (MLPs), subset frequencies were compared and revealed no significant differences between edited and control samples (Figure 4G; (47,49)). Additionally, editing efficiency was consistent across all populations, including LT-HSCs. These findings suggest that ABE-mediated disruption of ADGRE2 splice-sites is well-tolerated across all subsets of the hematopoietic hierarchy.

### *In vivo* pharmacology of ADGRE2 KO reveals no impact on engraftment, multilineage reconstitution, or editing persistence

To evaluate the long-term hematopoietic impact of ADGRE2 KO, we conducted a 16-week xenograft study using NOD-scid IL2Rγ^-^/^-^ (NSG) mice transplanted with ABE-edited (KO-ABE) or control human CD34^+^ HSPCs (Figure 5A). Assessment of BM samples revealed robust persistence of ADGRE2 splice-site disruption in all KO-ABE mice (median 98.4% editing efficiency versus 87.7% in input cells; Figure 5B). Human cell engraftment (chimerism) and multilineage reconstitution were comparable between KO-ABE and control groups (Figure 5C). Flow cytometry confirmed sustained and significant ADGRE2 surface loss across hematopoietic lineages in the KO-ABE group (Figure 5D). Similar trends were observed in peripheral blood (Supplemental Figures 3A, 3B). Together, these results demonstrate that ABE-mediated ADGRE2 KO supports long-term engraftment, preserves multilineage differentiation, and maintains stable editing, enabling effective target ablation without impairing hematopoietic function.

**Figure 5.**
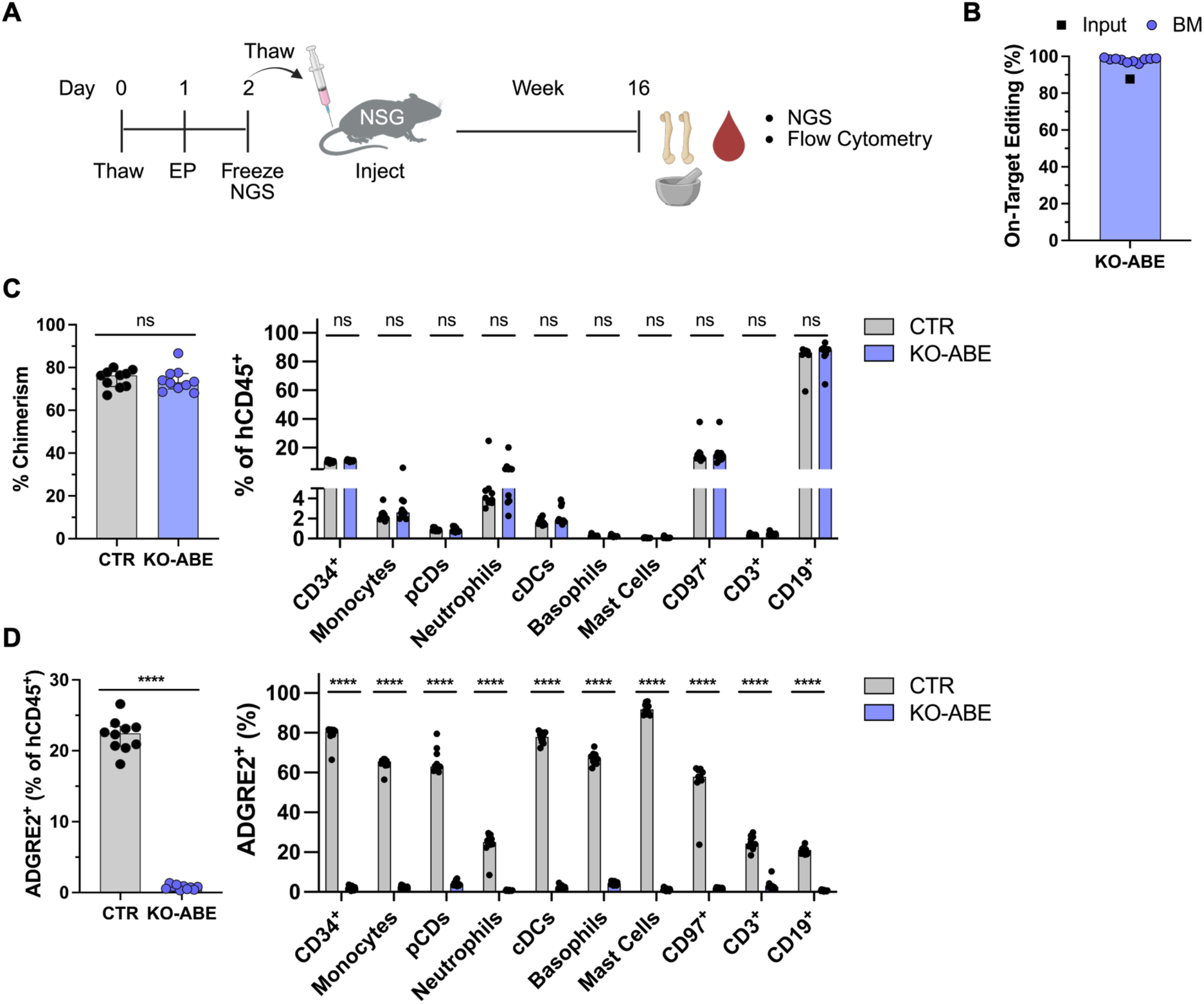
*In vivo* pharmacology study. **(A)** Experimental schema for 16-week xenotransplantation study in NSG mice. **(B)** KO-ABE and Control (CTR) HSPCs were frozen 48 h post-electroporation and then thawed for same-day injection into NSG mice (n=10 mice per group). Bone marrow (BM) was harvested after 16 weeks. On-target editing (A-to-G conversion at desired splice donor site) was assessed in input cells (pre-freeze) and BM samples by next-generation sequencing (rhAmpSeq). Black square indicates input material; each purple dot represents an individual mouse BM sample and horizontal bar denotes group mean. **(C)** Chimerism and multilineage reconstitution in BM, 16 weeks post-engraftment; measured by flow cytometry. Human chimerism calculated as hCD45^+^ / (hCD45^+^ + mCD45^+^) x 100. Frequencies of CD34^+^ HSPC, myeloid lineages (Monocytes, pDCs, Neutrophils, cDCs, Basophils, Mast Cells), and lymphoid (T cells, CD3^+^; B-cell, CD19^+^) populations were measured within hCD45^+^ compartment. CD97^+^ cells also assessed to confirm editing specificity. Data shown mean ± SD; n=10 mice per group. Each dot represents an individual mouse BM sample. Statistical analysis by two-way ANOVA; ns = not significant (p > 0.05). **(D)** ADGRE2 surface protein expression in total bone marrow hCD45^+^, myeloid, lymphoid, and CD97^+^ populations in KO-ABE versus CTR mice at study end; measured by flow cytometry. Data represented as mean ± SD; n=10 mice per arm, each dot represents one mouse. Statistical analysis performed using two-way ANOVA; ****p<0.0001. Abbreviations: NSG, NOD-scid IL2Rg null; KO, knockout; HSPC, hematopoietic stem and progenitor cells; hCD45, human CD45; mCD45, murine CD45; pDCs, plasmacytoid dendritic cells; cDCs, classical dendritic cells; SD, standard deviation.

## DISCUSSION

AML immunotherapy requires balancing therapeutic efficacy with the risks of antigen escape and on-target, off-tumor toxicity. The extensive biological heterogeneity of AML underscores the need to expand the repertoire of targetable antigens. In this study, we confirmed and extended prior reports of ADGRE2 expression in AML, showing consistent prevalence across blasts and LSCs, with surface intensities comparable to established AML targets such as CD123 and CLL-1, though lower than CD33 (19). Notably, ADGRE2 expression was observed on the highest percentage of cells among all markers on both blasts and LSCs irrespective of per cell expression, supporting its potential as a single or combinatorial immunotherapy target.

We generated and characterized a panel of ADGRE2-directed CAR T constructs, identifying ADGRE2-5-scFv as the lead candidate. This construct demonstrated potent, antigen-specific cytotoxicity against AML targets expressing even as few as ∼932 ADGRE2 surface antigens, a threshold within the range observed in primary AML cells. At a lower E:T of 1:4, ADGRE2-5-scFv CAR Ts exhibited robust cytotoxicity with minimal off-target activation, though other constructs demonstrated antigen-independent killing, emphasizing the need for rigorous binder selection. A notable observation was ADGRE2 upregulation on activated T cells, raising the possibility of fratricide or auto-activation, a challenge reminiscent of CD123-directed CAR therapies (37). Further studies using primary AML blasts of varying antigen levels, repeat challenge assays, and *in vivo* models will be critical to fully define the therapeutic threshold and development for clinical translation.

While the therapeutic potential of ADGRE2 is clear, the high expression in healthy HSPCs and mature myeloid lineages introduces a barrier to selective and safe targeting. In contrast to CD33, which is functionally dispensable (12,17,18,50), the biological necessity of ADGRE2 in the normal hematopoietic compartment and its myeloid function was previously unclear. Population-scale human genetics data provide evidence of *ADGRE2* dispensability. In gnomAD, nine distinct homozygous pLOF variants are carried by over 150 individuals, consistent with minimal evolutionary constraint (pLI = 0, LOEUF = 1.03). Functional characterization confirmed several *bona fide* KOs, including a truncating allele that reduced ADGRE2 surface expression by >85% in HSPCs from a single donor. While these findings provide genetic precedent for ADGRE2 dispensability, the current functional validation was performed in HSPCs from a single donor; additional evaluation across multiple donors will be required to confirm reproducibility.

To further interrogate the role of ADGRE2 in hematopoiesis beyond genetic association alone, we used both CRISPR-Cas9 and ABE-mediated strategies to disrupt *ADGRE2* and ablate surface expression in CD34^+^ HSPCs. Functional studies revealed that ADGRE2 deletion did not impair monocytic differentiation, viability, myeloid marker expression and did not alter the distribution of HSPC subsets, including LT-HSCs. Moreover, base editing at the exon 19 splice site resulted in efficient, precise, and durable gene disruption, leading to loss of ADGRE2 protein expression. Splice-site disruption was maintained across all HSPC subsets. *In vivo* pharmacology studies using NSG mice confirmed that ADGRE2-KO HSPCs retain the ability to engraft, support multilineage hematopoiesis, and maintain editing over a 16-week period. These findings mirror those seen with CD33-directed HSPC protection strategies (12,17,18) and support the use of similar immune-shielding approaches for ADGRE2-targeted therapies. Although long-term safety and persistence of *ADGRE2*-edited HSPCs—especially within the LT-HSC compartment—should be confirmed in serial transplantation studies.

Together, our findings establish ADGRE2 as both a biologically dispensable and therapeutically relevant AML target. The success of this approach would enable potent and selective ADGRE2-directed therapies as part of post-transplant relapse prevention or curative regimens by integrating protective genome-edited allogeneic donor-derived HSPC to mitigate toxicity. Future strategies may build on these findings by combining antigen ablation, epitope editing (51–53), and novel cell therapy platforms to safely broaden the therapeutic window in AML and other hematologic malignancies.

## Data availability statement

The publicly available human genetics data of ADGRE2 used in this study (related to Figure 3) are available in the gnomAD browser (https://gnomad.broadinstitute.org/gene/ENSG00000127507?dataset=gnomad_r4). All data generated or analyzed during this study are included in this published article and its supplementary information files. Source data are provided with this paper.

## Acknowledgments

We thank our colleagues at Vor Bio for their invaluable technical support and insightful contributions towards this study. We are especially grateful to the patients and healthy donors who generously provided samples for our research studies. We also acknowledge Ciara Tucker for facilitating cross-functional team communication and alignment and Ben Hall for his contributions to intellectual property drafting and publishing efforts. We acknowledge Abound Bio Inc. for developing the ADGRE2 CAR T binders and Siddhartha Mukherjee and Florence Borot for sharing the lead adenine base editor guide-RNA sequence, as well as their thoughtful scientific discussions and valuable input. Figures 1C, 3C, 3E, 4E, and 5A were created using BioRender.com. All *in vivo* experiments were conducted in accordance with relevant ethical regulations and standards for animal testing. Protocols were approved by the Institutional Animal Care and Use Committee (IACUC).

## Author Contributions

Y.K., J.E., M.S., H.M., H.H., J.S., and J.R.L conceived and designed the study. Y.K., J.E., M.S, H.M., H.H., N.M, and A.H. performed the experiments. Data was analyzed by Y.K., H.M, H.H, N.M., A.H., H.Q., and R.W., with input from J.E., M.S., J.X.F., M.I.L, J.S., and J.R.L. Statistical analyses were carried out by Y.K., J.E., and N.M.. The manuscript was drafted by Y.K., J.E., M.S., J.S., and J.R.L.. All authors critically revised the manuscript for intellectual content. J.E., M.S., J.X.F., M.I.L., H.G.G., J.S., T.C., and J.R.L. supervised the study and coordinated collaboration across the research teams. All authors approved the final version of the manuscript.

