## Supplemental Information for "Integrating Human Genetics and Protective Genome Editing to Enable ADGRE2-Directed AML Therapy"

**Title:**

[1] Vor Bio, Cambridge, MA, 02140, USA.

[2] Current Address: National Institute of Mental Health (NIMH), Bethesda, MD, 20892, USA.

[3] Current Address: UMass Chan Medical School, Worcester, MA, 01655, USA.

### Supplementary Materials

#### *Primary AML patient samples*

Patient details can be found in Supplemental Table 1. All samples were de-identified and collected under the approved protocols by the Independent Ethics Committee of Arte Med Assistance, LLC with informed patient consent in accordance with the Declaration of Helsinki. Data acquisition was performed on the Cytek® Aurora (Cytek Biosciences, Fremont, CA, USA). A complete list of antibodies is provided in Supplementary Table 2. Five samples were excluded from LSC analysis due to low (<100) cell count.

#### *Healthy donor samples*

Samples were received fresh and red blood cell lysis was performed by adding ammonium chloride solution (STEMCELL Technologies Inc., Cambridge, MA, USA). A complete list of antibodies and cell type gating strategy is provided in Supplemental Tables 3 and 4. Fluorescence-minus-one (FMO) controls were used to determine marker positivity.

#### *Primary human CD34+ cells*

Cryopreserved CD34+ isolated cells from the peripheral blood of dual mobilized (G-CSF + Plerixafor) healthy donors were received from Charles River Laboratories (Wilmington, MA, USA) or Stem Express. Cells were thawed and washed with DPBS + 10% HSA and maintained in StemSpan SFEM (STEMCELL Technologies) or CellGenix® SCGM (Sartorious, Cambridge,

MA, USA) media formulated with stem cell factor (SCF), Fms-related tyrosine kinase 3 ligand (FLT-3L), and thrombopoietin (TPO) at 37°C, 5% $CO_2$ and 95% humidity.

***Cell Lines***

MOLM-13 (AcceGen, Fairfield, NJ, USA) and Jurkat (ATCC, Manassas, VA, USA) cell lines were cultured in RPMI-1640 (Thermo Fisher Scientific, Waltham, MA, USA) supplemented with 10% heat-inactivated FBS (Corning, Tewksbury, MA, USA). HL-60 (ATCC) cell line was cultured in IMDM (Thermo Fisher Scientific) supplemented with 20% heat-inactivated FBS (Corning). HEK293T (ATCC) cells were cultured in DMEM (Thermo Fisher Scientific) supplemented with 10% heat-inactivated FBS (Corning) and sub-cultured per recommended protocols using 0.25% Trypsin-EDTA for trypsinization. Suspension-adapted HEK293F (Themo Fisher Scientific) were cultured. Suspension-adapted HEK293F were used for lentivirus production and were acquired from ThermoFisher. All cell lines were maintained at 37°C, 5% $CO_2$ and 95% humidity.

***Primary T-cells, activation, and culture***

Cryopreserved human peripheral blood $CD3^+$ pan-T cells (sourced from AllCells, Alameda, CA, USA or STEMCELL Technologies) were thawed in 100% FBS and cultured in Optimizer media supplemented with 2.6% CTS Optimizer supplement, 2.5% CTS Immune Cell SR, 1% L-Glutamine, 1% Glutamax (from Thermo Fisher Scientific), and 1% rh IL-2 (Cellgenix; 100,000 u/mL stock,). After 24 h, T cells were activated with human CD3/CD28 TransAct beads (Miltenyi Biotec, Charlestown, MA, USA) at a 1:100 dilution in complete T-cell media. Two days post-activation, cells were harvested for flow cytometry to assess ADGRE2 expression on the Novocyte Quanteon (Agilent Technologies, Santa Clara, CA, USA). Non-activated T cells were cultured in parallel. Markers included in the flow cytometry panel were Fixable Viability Stain 780 (BD Biosciences, Franklin Lakes, NJ, USA), EMR2 2A1 PE (Miltenyi Biotec), and CD3 BV605, CD4 BV421, CD8 BV510, CD69 AF700, and CD25 PerCP/Cy5.5 from BioLegend (San Diego, CA, USA). Catalog numbers can be found in Supplemental Table 5.

***Construction of expression vectors***

For CAR binding studies, expression DNA constructs of ADGRE2$^{\text{EGF1,2,3,4,5}}$ wild-type coding sequence (CDS) (CCDS32935.1) or its modified isoforms were cloned into the pcDNA3.1(+)-P2A-eGFP backbone vector (GenScript). Isoform-specific constructs included ADGRE2$^{\text{EGF1,2,3,5}}$ (deletion of CDS sequence for exon 7; nucleotides 488-634), ADGRE2$^{\text{EGF1,2,5}}$ (deletion of CDS sequence for exons 6 and 7; nt 356-634), and ADGRE2$^{\text{EGF1,2}}$ (deletion of CDS sequence for exons 6, 7 and 8; nt 356-781). ADGRE2$^{\text{GAIN+GPS repl.}}$ had replacement of ADGRE2 CDS nt 874-1590 with the corresponding region from CD97 (ADGRE5; CCDS32929.1; nt 865-1626). In all ADGRE2 constructs, an HA tag (YPYDVPDYA) CDS was inserted directly downstream of the ADGRE2 signal peptide (nt 1-69). The full-length (FL) CD97 (ADGRE5) was generated from CCDS32929.1, with the HA tag inserted directly downstream of its signal peptide (nt 1-60).

For studies of naturally occurring ADGRE2 variants, constructs were cloned into the pcDNA3.1(+)-IRES-eGFP backbone vector (GenScript) based on the WT ADGRE2 sequence (CCDS32935.1). Point mutations and deletions corresponding to single-point mutations (SNPs) reported in gnomAD v4.1.0 were introduced as follows: Variant A (rs143325445) c.436 C>T, Variant B (rs1183101283) c.370 C>T, Variant C (rs559663437) c.396 deletion (del.), Variant D (rs1401601834) c.838 C>T, Variant E (rs754549569) c.1581 C>A, Variant F (rs534018804) c.1755_1756 del., Variant G (rs375427337) c.1693_1709 del., Variant H (rs548759776) c.2124 G>A. Each construct also included a FLAG tag (DYKDDDDK) CDS inserted immediately downstream of the ADGRE2 signal peptide (nt 1-69).

### Supplementary Methods

#### *Flow Cytometry*

Cells were collected and washed with FACs buffer containing 1% bovine serum albumin (BSA; Rockland Immunochemicals, Pottstown, PA, USA). For samples derived from primary human sources, Fc receptors were blocked using human FcX TruStain and True-Stain Monocyte Blocker (BioLegend), each diluted at 1:20 in FACs buffer. For xenotransplant mouse samples, TruStain FcX PLUS anti-mouse CD16/32 (BioLegend, Catalog # 156604) was also included in the blocking mixture at a 1:20 dilution. Following blocking, cells were stained with fluorophore-conjugated antibodies at room temperature for 20 minutes or on ice for 30 minutes. Samples were acquired on one of the following instruments: Novocyte Quanteon (4 laser-configuration; Agilent Technologies), CytoFlex LX flow cytometer (6 laser-configuration; Beckman Coulter, Brea, CA, USA), or Aurora Spectral Flow Cytometer (5-laser configuration; Cytek Biosciences). Data was analyzed using FlowJo software (BD Biosciences). Data visualizations and statistical analyses were generated and completed with GraphPad Prism software for MacOS (San Diego, CA, USA).

***DNA Plasmid Transfections***

HEK293T cells were seeded at 8 x $10^5$ cells per well of 6-well plates using DMEM supplemented with 10% heat-inactivated FBS. After 24 h, cells were transfected with 2 μg of DNA plasmid (GenScript, Piscataway, NJ, USA) per well. DNA was pre-incubated with Lipofectamine 2000 reagent (Thermo Fisher Scientific) at a 1:2 ratio in Opti-MEM for 10 minutes at room temperature and then added dropwise to the culture wells and gently mixed using figure-eight motion. Following transfection, cells were incubated for 24 hours at 37°C, 5% $CO_2$ and 95% humidity before trypsinization and collection for downstream analysis.

***ADGRE2 CAR antibody construction and detection***

ADGRE2-directed CAR antibodies were produced by Abound Bio Inc. (Pittsburgh, PA, USA) and supplied at a concentration of 250nM, with a C-terminal FLAG (DYKDDDDK) peptide tag for detection. For flow cytometry assays, single-chain variable fragment (scFv) and heavy chain variable domain (VH) CAR formats were used at a final concentration of 6.25μg/mL and 3.75μg/mL per test, respectively. Detection of CAR antibodies was performed using anti-FLAG Clone L5 APC antibody (BioLegend, Catalog #637308) as a secondary reagent. Samples were collected on the Novocyte Quanteon (Agilent Technologies), and data was analyzed using FlowJo software. Dead cells were excluded using DAPI (4,6-Diamidino-2-Phenylindole; BioLegend, Catalog #422801) and subsequent analysis was restricted to transfected cells ($eGFP^+$).

***Virus and CAR T cell production***

CAR containing plasmids (GenScript) were transfected into suspension-adapted HEK293F cells cultured in LV-Max Production Medium using the LV-MAX Transfection Kit (all from Thermo Fisher Scientific) as per manufacturer's instructions. Briefly, transfection complexes were prepared by combining 15µL of LV-Max Transfection Reagent with 10µg of plasmid DNA, followed by the addition of 60µL of LV-MAX Transduction Reagent (diluted in Opti-MEM). Complexes were incubated at room temperature for 10 minutes before being added (1mL) to the HEK293F cells. Cells were incubated at 37°C, 8% $CO_2$ on an orbital shaker (240 rpm). LV-Max Enhancer (400µL) was added after 5-6 hours and then returned to the incubator. After a total incubation time of 24-hours, viral supernatant was collected via ultracentrifugation using an Optima XE-90 Ultracentrifuge (Beckman Coulter), resuspended in PBS, and stored at -80°C. Pan-T cells were thawed and underwent same day human CD4/CD8 positive isolation via MACs magnetic separation (Miltenyi Biotec) and activation with human CD3/CD28 TransAct beads (1:16.51 TransAct:Media; Miltenyi Biotec, Catalog # 130-111-160). After 48 hours, activated T cells underwent lentivirus transduction using the TransDux MAX Virus Transduction Reagent Kit (SBI System Biosciences, Palo Alto, CA, USA) following manufacturer instructions. Transduced cells were cultured in 6-well G-Rex plates (Wilson Wolf Manufacturing, St Paul, MN, USA) in complete T cell optimizer media. Untransduced T-cells were cultured in separate wells as controls. Cells were incubated at 37°C, 5% $CO_2$, and 95% humidity. Additional T-cell media was added after 24 hours (100mL total), and cells were harvested after an additional 4 or 5 days for viability assessment and CAR detection by flow cytometry. All remaining cells were viably cryopreserved in CryoStor® 10 (BioLife Solutions, Inc., Bothell, WA, USA) and stored at $LN_2$.

***In vitro cytotoxicity assays***

*In vitro* cytotoxicity assays were conducted using ADGRE2-directed CAR T cells against MOLM-13 target cell lines to assess CAR T cell efficacy. ADGRE2-knockout (KO) MOLM-13 were generated via CRISPR-Cas9-mediated gene editing, followed by single-cell sorting and expansion of a confirmed ADGRE2-null clone. Effectors (E) and CFSE-labeled Targets (T) were co-cultured in triplicate at a 1:1 ($5 \times 10^4$ targets and $5 \times 10^4$ $CAR^+$ effectors) or 1:4 E:T ratio in 96-well flat-bottom plates. Co-cultures were incubated at 37°C, 5% $CO_2$ and 95% humidity for 24 or 48 hours in MOLM-13 complete medium (200µL/well). Control wells containing only effectors or targets were plated in replicates as controls. Following incubation, cells were harvested for flow cytometry, analyzed, and compensated on the Novocyte Quanteon (Agilent). Viable target cells were identified as Annexin-$V^-$/LIVE-$DEAD^-$. CAR T cells were detected using AffiniPure Goat Anti-Human IgG (H+L) (Jackson ImmunoResearch, West Grove, PA, USA), which was applied prior to Fc receptor blocking (FcX and monocyte receptor blockade). A complete list of antibodies is provided in Supplemental Table 5. Additionally, supernatants were collected for cytokine profiling using the Human CD8+ T-Cell Magnetic Bead Pre-Mixed 17-Plex kit (Millipore Sigma, Darmstadt, Germany), with data acquisition on the Luminex FlexMap 3D system (Luminex, Austin, TX, USA).

***Engineered Antigen Diverse Clones***

MOLM-13 AML cells were engineered to stably express a range of variable but distinct ADGRE2 surface antigen molecules. Lentiviral vectors (LVV) were constructed by cloning the

ADGRE2 coding sequence (CCDS32935.1) into the pGenLenti backbone (GenScript) under a CMV or UBC promoter. DNA plasmids were transfected into suspension-adapted HEK293F cells and 50uL of fresh virus was transduced into ADGRE2-KO MOLM-13 cells (CRISPR-Cas9 generated followed by single cell sort; also KO for CD33 and CLL1) similar to CAR T transduction protocol above. After five days of incubation at 37°C, 5% $CO_2$, 95% humidity, culture media were replaced. Cells underwent single-cell sorting (BD FACSAria Fusion) into 96-well round-bottom plates containing 100µL of MOLM-13 complete medium. Clonal populations were expanded and ADGRE2 expression was assessed via flow cytometry.

***Western blotting***

HEK293T cells were scraped from culture wells and lysed in 4x Laemmli Sample Buffer (Bio-Rad) containing 10% 2-Mercapthoethanol (Bio-Rad). Lysates were boiled at 100°C for 2-5 minutes and centrifuged at maximum speed for 15 minutes at 4°C. Supernatants were collected and 20µL of sample was loaded onto Mini Protean TGX Precast Gels 4-20% precast polyacrylamide gels (12-well format; Bio-Rad). Precision Plus Protein™ All Blue Standards (Bio-Rad) were loaded into flanking lanes. Electrophoresis was performed at 100 V for 10 minutes followed by 160 V for ~80 minutes in 1x Tris/Glycine/SDS running buffer (10x stock, Bio-Rad). Transfer buffer was prepared using 1x Tris/Glycine buffer (10x stock, Bio-Rad) with 20% methanol (Sigma-Aldrich, St. Louis, MO, USA). Immobilon-FL PVDF membranes (Millipore) were activated in methanol (Millipore Sigma) for 2 minutes before equilibration in transfer buffer with mini trans-blot filter paper (Bio-Rad). Transfer sandwiches using foam pads (Bio-Rad) were assembled, and proteins were transferred at 120 V for 90 minutes in a cold environment (ice bucket with cold packs). Membranes were blocked in Intercept® Protein-Free

Blocking Buffer with TBS (LI-COR, Lincoln, NE, USA) for 30 minutes at room temperature, then incubated overnight at 4℃ with primary antibodies diluted in 1x TBS (Bio-Rad) + 0.1% Tween-20 (Bio-Rad). Primary antibodies used were Mouse anti-human EMR2 (clone 2A1; Invitrogen, Catalog #MA5-28205) at 1:1000, Purified anti-HA.11 (clone 16B12; BioLegend, Catalog # 901501) at 1:500, and Rabbit anti-β-actin (clone D6A8; Cell Signaling Technology, Catalog # 8457S) at 1:1000 in final solution. Following washes with 1x TBS with 0.1% Tween 20, membranes were incubated with secondary antibodies diluted in 1x TBS + 0.1% Tween-20 for 1 hour at room temperature. Secondary antibodies used were IRDye® 800CW goat anti-rabbit IgG (LICOR, Catalog # 926-32211) and IRDye 680RD goat anti-mouse IgG (LICOR, Catalog # 926-68070) each at a final 1:10,000 dilution. After additional washes with 1x TBS + 0.1% Tween-20 and final rinses with 1x TBS, membranes were imaged using the Bio-Rad ChemiDoc™ MP Imaging System.

***Transcript Expression***

ADGRE2 transcript levels were quantified using digital droplet PCR (ddPCR) methods (Bio-Rad, Hercules, CA, USA). Total RNA was isolated using the RNeasy Mini Kit (Qiagen, Germantown, MD, USA) following the manufacturer's instructions. RNA concentration was measured with the Qubit RNA BR Kit (Thermo Fisher Scientific). Subsequently, 100ng of RNA was reverse transcribed into cDNA using the QuantiTect Reverse Transcription Kit (Qiagen), according to the manufacturer's protocol. ddPCR reactions were prepared using 3ng cDNA, 10µL of 2x ddPCR Supermix for Probes (Bio-Rad), and 0.005nmol probe/0.01nmol primer (Integrated DNA Technologies (IDT), Coralville, IA, USA) of primer per reaction. Duplex reactions were prepared using custom ADGRE2 assays (6-FAM) and the GUSB reference gene assay (SUN; Hs.PT.58v.27737538, IDT). Custom ADGRE2 primer-probe sets (IDT PrimerQuest Tool) are as follows: Exons 10-12 (Primer 1 *CAGGCTTGGCCAATAACAC*, Primer 2 *GTCTACTTGCTTCTGCACCTC,* Probe /56-FAM/*TCCTCAGAG/ZEN/GCCTGAGCAAGAACC/3IABkFQ/*) and Exon 6 (Primer 1 *GCTTCGGGTCCTCAGGTTTGAG*, Primer 2 *GTCAGCAGAACCCAAGGCTCTG*, Probe /56-FAM/*TGTAGCTGC/ZEN/CGAGGGTGTTGAC/eIABkFQ/*). Reactions were run on the Bio-Rad QX200 ddPCR system. Droplet data were analyzed using Bio-Rad QX Manager Software to determine the number of positive droplets per sample. ADGRE2 transcript expression was normalized to GUSB and reported as: Normalized Expression = (ADGRE2 copies per cell / GUSB copies per cell) x 100.

***ADGRE2 Gene Editing in Human CD34+ Cells***

ADGRE2 gene disruption was achieved in $CD34^+$ HSPCs with either CRISPR-Cas9, adenine base editing (ABE), or cytosine BE (CBE) to induce protein KO or disruption via electroporation (EP). Lead guide-RNAs (gRNA) were identified after evaluation of editing efficiency and surface protein reduction. Prior to electroporation (EP), HSPCs were thawed and cultured for two days. For CRISPR-Cas9 mediated KO, the sNLS-SpCas9-sNLS nuclease (Aldevron, South Fargo, ND, USA; Catalog # 9212-0.25mg) protein and gRNA (Axolabs) were co-delivered into the cells as a pre-complexed ribonucleoprotein (RNP). For ABE or CBE editing, gRNA (Axolabs or Synthego, Redwood, CA, USA) was added to cells, followed by the addition of ABE8.20-m, BE4-PpAPOBEC1, or BE4-PpAPOBEC-SpG mRNA (GenScript) immediately before EP. All EPs were performed using either the Lonza Nucleofector system (1-2 x $10^6$ cells per EP, P3 Primary Cell 4D-Nuceleofector, CA-137 program; Lonza, Walkersville, MD, USA) or the MaxCyte GTx platform (10-12 x $10^6$ cells per EP, HyClone MaxCyte electroporation buffer (MxCEB), HSC3 program; Maxcyte, Inc., Rockville, MD, USA) following the manufacturer's recommended protocols. Post-EP, cells were cultured at 0.5-1 x $10^6$ cells/mL in HSPC culture media and incubated at 37C, 5% $CO_2$ and 95% humidity. The gRNA sequences used for KO were: CRISPR-Cas9 gRNA CTTGGCCAATAACACCATCC (PAM: AGG) and ABE8.20-m gRNA (generously provided by Dr. Mukherjee's lab) GTGGTACCTGCTGGCTGAGG (PAM: AGG). Further details on these guides can be found in the associated patent WO2023086422. The gRNA sequences used to induce variant D and H mutations were CCGATTCTTCGACAAAGTCC (PAM: AGG) and AATCCAGAGAGTCACCAGAA (PAM: AGA), respectively.

***Analysis of HSPC sub-populations***

Human $CD34^+$ HSPCs were electroporated (EP) with the MaxCyte GTx platform. After 48 hours post-EP, cells were stained with flow cytometry techniques and sorted into defined subpopulations using either the BD FACSAria Fusion (BD Biosciences) or SONY MA900 (Sony Biotechnology, San Jose, CA, USA) cell sorters. Cells were gated into the following subpopulations based on well-established surface marker profiles (1–4): common myeloid progenitors (CMPs), multi-lymphoid progenitors (MLPs), multipotent progenitors (MPPs), and long-term hematopoietic stem cells (LT-HSCs). Subpopulation frequencies were calculated as a percentage of total viable cells. A complete list of antibodies can be found in Supplemental Table 7. Gating was performed using fluorescence-minus-one (FMO) controls. The following gating strategies were used to define subpopulations: CMPs ($CD34^+CD38^+CD45RA^-$), MLPs

($CD34^+CD38^-CD45RA^+CD90^-$), MPPs ($CD34^+CD38^-CD45RA^-CD90^-$), LT-HSCs ($CD34^+CD90^+CD45RA^-EPCR^+ITGA3^+CD133^+$).

***In vivo pharmacology***

*In vivo* pharmacology studies were performed using a xenotransplant NOD-scid IL2Rg null (NSG) mouse model to evaluate long-term engraftment, hematopoietic reconstitution across multiple lineages, and persistence of ADGRE2 gene editing. ADGRE2-KO samples were generated via adenine base editing (ABE8.20-m) using the MaxCyte GTx device (Maxcyte, Inc., Rockville, MD, USA). Forty-eight hours post-electroporation edited and non-edited (control) $CD34^+$ HSPCs were cryopreserved in CryStor® 10 (Biolife Solutions). On the day of engraftment, cells were thawed and administered intravenously into sublethally irradiated NSG mice. Engraftment and multilineage hematopoietic reconstitution were assessed 16 weeks post-engraftment. Peripheral blood and bone marrow were analyzed by flow cytometry using human and murine CD45 markers, along with lineage-specific markers for human hematopoietic cell populations and ADGRE2 surface expression (see Supplemental Table 8 for antibody panel). Persistence of ADGRE2 editing was assessed in engrafted human cells by isolating genomic DNA followed by next-generation sequencing.

***CRISPR-Cas9 gene editing frequency by Sanger sequencing***

Genomic DNA (gDNA) was isolated from edited and control cells using either the ZyGEM prepGEM kit or DNA Blood & Tissue Kit (Qiagen), following the manufacturer's protocols. PCR amplification was performed using Q5 High Fidelity 2X Master Mix (New England

Biolabs, Ipswich, MA, USA) and custom-designed primers (10uM each; IDT). Primer sequences were as follows: Forward CTCAGAGGCCAACACTTCTGTGC, Reverse TCCACCGTCTGCTTCAACACCG. PCR products were purified using either the QIAquick PCR Purification Kit (Qiagen) or AMPure XP beads (Beckman Coulter). For bead-based purification, PCR products were incubated with AMPure XP beads for 5-10 minutes at room temperature. After magnetic separation, the supernatant was discarded, and the beads were washed twice with 70% ethanol. Beads were then air-dried on the magnetic plates and eluted in nuclease-free water. The purified PCR product was quantified from the supernatant following final magnetic separation. Sanger sequencing was performed by Azenta Life Sciences (Burlington, MA, USA) using the sequencing primer CCTAGAAGCTCCCCACC. Editing frequency and insertion and deletion (indel) profiles were analyzed using the Interference of CRISPR Edits (ICE) analysis platform (5).

***ABE gene editing frequency by Next Generation Sequencing***

Genomic DNA was extracted as previously described. On-target editing resulting from ABE8.20-m base-editing was evaluated by next generation sequencing (NGS) using rhAmpSeq (IDT) technology. PCR primers were designed with the rhAmpSeq design tool to amplify regions surrounding the target site (insert size: 150-350 bp) with overhang Illumina (Illumina, San Diego, CA, USA) adapters. Sample libraries were barcoded, prepared according to the manufacturer's instructions, and sequenced using the Miseq Sequencer System (Illumina). Denatured PhiX Control v3 (Illumina) was added to the pooled library at 1:10 (PhiX:sample) volume ratio to improve sequence diversity and quality control. For data processing and analysis, mutations within the amplified regions were identified and quantified using CRISPRessoWGS from the CRISPResso analysis suite (6). The quantification window was set to include 10 bp flanking both sides of the target nucleotide (quantification_window_size = 10). Variant consequences were annotated using the Ensembl Variant Effect Predictor (VEP; (7)) in the offline mode. Customized high-level functional classifications were assigned to each mutation based on its predicted consequences: Stop Gained—contains 'stop_gained', InDels—insertions or deletion in the alignment, Splice Site Disruption—contains 'splice_', Missense Variants—all other substitutions not meeting the above criteria. If a mutation matched multiple categories, classification was prioritized in the following hierarchy: Stop Gained > InDels > Splice Site Disruption > Missense Variants. The frequency of each functional class was calculated by summing the percentages of all associated mutations.

**Supplemental Table 1. AML Patient Characteristics**

| **Characteristic** | N=25[1] |
|---|---|
| **Disease Status** | |
| Diagnosis | 23 (92%) |
| Transformation from MDS | 3 (13%) |
| Relapse | 2 (16%) |
| **Sex** | |
| Female | 15 (60%) |
| Male | 10 (40%) |
| **Age at Diagnosis** | 68 (56-71) |
| **Blast %** | |
| at Diagnosis | 70.4 (41.5-80) |
| at Relapse | 94.3 (92-96) |
| **Previous Treatment** | 4 (16%) |

[1] n (%); Median (Q1, Q3)

**Supplemental Table 2**.

| **Reagent** | **Vendor** | **Catalog Number** |
|---|---|---|
| DAPI 4,6-Diamidino-2-Phenylindole | BioLegend | 422801 |
| anti-human Lineage Cocktail BV510™ (CD3 OKT3, CD14 M5E2, CD16 3G8, CD19 | BioLegend | 348807 |

| | | |
|---|---|---|
| HIB19, CD20 2H7, CD56 HCD56) | | |
| anti-human CD45 Alexa Fluor® 488 (clone HI30) | BioLegend | 304017 |
| anti-human CD34 APC/Fire™ (clone 581) | BioLegend | 343536 |
| anti-human CD38 BUV395™ (clone HB7) | BD Biosciences | 563811 |
| anti-human CD312 (EMR2) PE (clone REA302 (2A1)) | Miltenyi Biotec | 130-119-770 |

**Supplemental Table 3**.

| Reagent | Vendor | Catalog Number |
|---|---|---|
| LIVE/DEAD™ Fixable Blue Dead Cell Stain, for UV excitation | Invitrogen | L34962 |
| anti-human CD45 BUV395™ (clone HI30) | BD Biosciences | 563792 |
| anti-human CD312 (EMR2) PE (clone REA302 (2A1)) | Miltenyi Biotec | 130-119-770 |
| Anti-human CD45RA FITC (clone HI100) | BioLegend | 304106 |
| anti-human CD90 (Thy1) PE/Dazzle™ 594 (clone 5E10) | BioLegend | 328134 |
| anti-human CD38 BV510™ (clone HIT2) | BioLegend | 303540 |
| Anti-human CD10 APC (clone HI10a) | BioLegend | 982202 |
| Anti-human CD135 BV786™ (clone 4G8) | BD Biosciences | |

| | | |
|---|---|---|
| Human Hematopoietic Lineage Antibody, eFluor™ 450 (CD2 RPA-2.10, CD3 OKT3, CD14 61D3, CD16 CB16, CD19 HIB19, CD56 TULY56, CD235a HIR2) | eBioscience™ (Thermo Fisher) | 22-7775-72 |
| Anti-human CD14 BV711™ (clone M5E2) | BioLegend | 301838 |
| Anti-human CD16 FITC (clone 3G8) | BioLegend | 302006 |
| Anti-human CD117 (c-kit) APC/Cyanine7 (clone 104D2) | BioLegend | 313228 |
| Anti-human CD203c (E-NPP3) APC (clone NP4D6) | BioLegend | 324610 |
| Anti-human CD303 (BDCA-2) BV421™ (clone 201A) | BioLegend | 354212 |
| Anti-human CD3 BV605™ (clone OKT3) | BioLegend | 317322 |
| Anti-human HLA-DR BV510™ (clone G46-6) | BD Biosciences | 563083 |
| Anti-human CD11c PerCP-Cy™5.5 (clone B-ly6) | BD Biosciences | 656145 |
| Anti-human CD125 BV786™ (clone A14) | BD Biosciences | 743932 |
| Anti-human CD19 Alexa Fluor® 700 (clone HIB19) | BD Biosciences | 561031 |
| Anti-human CD56 BUV737™ (clone MY31) | BD Biosciences | 748609 |

**Supplemental Table 4.**

| Population | Gating Strategy |
|---|---|
| HSC | hCD45+Lin-CD34+CD38-CD45Ra-CD90+ |
| MPP | hCD45+Lin-CD34+CD38-CD45Ra-CD90- |
| CMP | hCD45+Lin-CD34+CD38+CD10-CD45Ra-CD135+ |
| GMP | hCD45+Lin-CD34+CD38+CD10-CD45Ra+CD13+ |
| MEP | hCD45+Lin-CD34+CD38+CD10-CD45Ra-CD135- |
| CLP | hCD45+Lin-CD34+CD38+CD10+ |
| Basophils | hCD45+CD3-CD19-CD56-HLA-DR-CD16-CD14-CD203c+CD117- |
| Eosinophils | hCD45+CD16-SSChi |
| Neutrophils | hCD45+CD16+SSChi |
| cDCs | hCD45+HLA-DR+CD14-CD303-CD11c+ |
| pDCs | hCD45+HLA-DR+CD14-CD303+CD11c- |
| Monocytes | hCD45+FSCmedSSCmedCD14+CD16- |
| Mast cells | hCD45+CD3-CD19-CD56-HLA-DR-CD16-CD14-CD203c+CD117+ |
| T cells | hCD45+FSCloSSCloCD3+ |
| B cells | hCD45+FSCloSSCloCD19+ |
| NK cells | hCD45+FSCloSSCloCD56+ |

344

**Supplemental Table 5.**

| Reagent | Vendor | Catalog Number |
|---|---|---|
| Fixable Viability Stain 780 | BD Biosciences | 565388 |
| Anti-human CD312 (EMR2) PE (clone REA302 (2A1)) | Miltenyi Biotec | 130-119-770 |
| PE Annexin V | BioLegend | 640908 |
| Anti-human CD3 BV605™ (clone UCHT1) | BioLegend | 300460 |
| Anti-human CD4 BV421™ (clone RPA-T4) | BioLegend | 300532 |
| Anti-human CD8 BV510™ (clone RPA-T8) | BioLegend | 301048 |
| Anti-human CD25 PerCP/Cyanine5.5 (clone BC96) | BioLegend | 302626 |
| Anti-human CD69 Alexa Fluor® 700 (clone FN50) | BioLegend | 310922 |
| Alexa Fluor® 647 AffiniPure Goat Anti-Human IgG (H+L) | Jackson ImmunoResearch | 109-605-003 |

**Supplemental Table 6.**

| Reagent | Vendor | Catalog Number |
|---|---|---|
| LIVE/DEAD™ Fixable Blue Dead Cell Stain, for UV excitation | Invitrogen | L34962 |
| Anti-human CD11b APC/Cyanine7 (clone ICRF44) | BioLegend | 301342 |
| Anti-human CD33 BV711™ (clone P67.6) | BioLegend | 366624 |

| | | |
|---|---|---|
| Anti-human CD14 BV421™ (clone M5E2) | BioLegend | 301830 |
| Anti-human CD15 BB515™ (clone HI98) | BD Biosciences | 565236 |

**Supplemental Table 7.**

| **Reagent** | **Vendor** | **Catalog Number** |
|---|---|---|
| LIVE/DEAD™ Fixable Aqua Dead Cell Stain, for 405nm excitation | Invitrogen | L34966 |
| Anti-human CD34 FITC (clone 581) | BD Biosciences | 555821 |
| Anti-human CD201 (EPCR) PE (clone RCR-401) | BioLegend | 351904 |
| Anti-human CD90 (Thy1) APC/Cyanine7 (clone 5E10) | BioLegend | 328132 |
| Anti-human CD90 (Thy1) PE-Dazzle™ 594 (clone 5E10) | BioLegend | 328134 |
| Anti-human CD133 BB700™ (clone 293C3) | BD Biosciences | 747563 |
| Anti-human CD49c BV605™ (clone C3 II.1) | BD Biosciences | 744518 |
| Anti-human CD45RA BV711™ (clone HI100) | BioLegend | 304137 |
| Anti-human CD10 BV421™ (clone HI10a) | BioLegend | 312218 |
| Anti-human CD38 Alexa Fluor® 657 (clone HIT2) | BioLegend | 303514 |

**Supplemental Table 8.**

| Reagent | Vendor | Catalog Number |
| --- | --- | --- |
| LIVE/DEAD™ Fixable Near-IR (876) Dead Cell Stain, for 808 nm excitation | Invitrogen | L34982 |
| anti-hu CD312 (EMR2) PE (clone REA302 (2A1)) | Miltenyi Biotec | 130-119-770 |
| anti-human CD45 BUV395™ (clone HI30) | BD Biosciences | 563792 |
| Anti-mouse CD45 BV650™ (clone 30-F11) | BioLegend | 103151 |
| Anti-human CD3 APC/Cyanine7 (clone OKT3) | BioLegend | 317342 |
| anti-human CD19 BB515™ (clone HIB19) | BD Biosciences | 564456 |
| anti-human CD33 PE/Cyanine7 (clone P67.6) | BioLegend | 366618 |
| Anti-human CD34 BV786™ (clone 581) | BD Biosciences | 743534 |
| Anti-human CD15 (SSEA-1) Alexa Fluor® 700 (clone HI98) | BioLegend | 301920 |
| Anti-human CD14 BUV661™ (clone M5E2) | BD Biosciences | 741603 |
| Anti-human HLA-DR BV510™ (clone L243) | BD Biosciences | 563083 |
| Anti-human CD203c (E-NPP3) BV605™ (clone NP4D6) | BioLegend | 324620 |
| Anti-human CD11c BV421™ (clone B-ly6) | BD Biosciences | 562561 |

| | | |
|---|---|---|
| Anti-human CD303 (BDCA-2) PE-Vio® 615 (clone AC144) | Miltenyi Biotec | 130-113-752 |
| Anti-human CD97 APC (clone 380903) | R&D Systems | FAB2529A |
| Anti-human CD10 BUV496™ (clone HI10a) | BD Biosciences | 741137 |
| Anti-human CD117 (c-kit) PE/Cyanine5 (clone 104D2) | BioLegend | 313210 |
| Anti-human CD33 PerCP/Cyanine5.5 (clone HIB22) | BioLegend | 302516 |

352

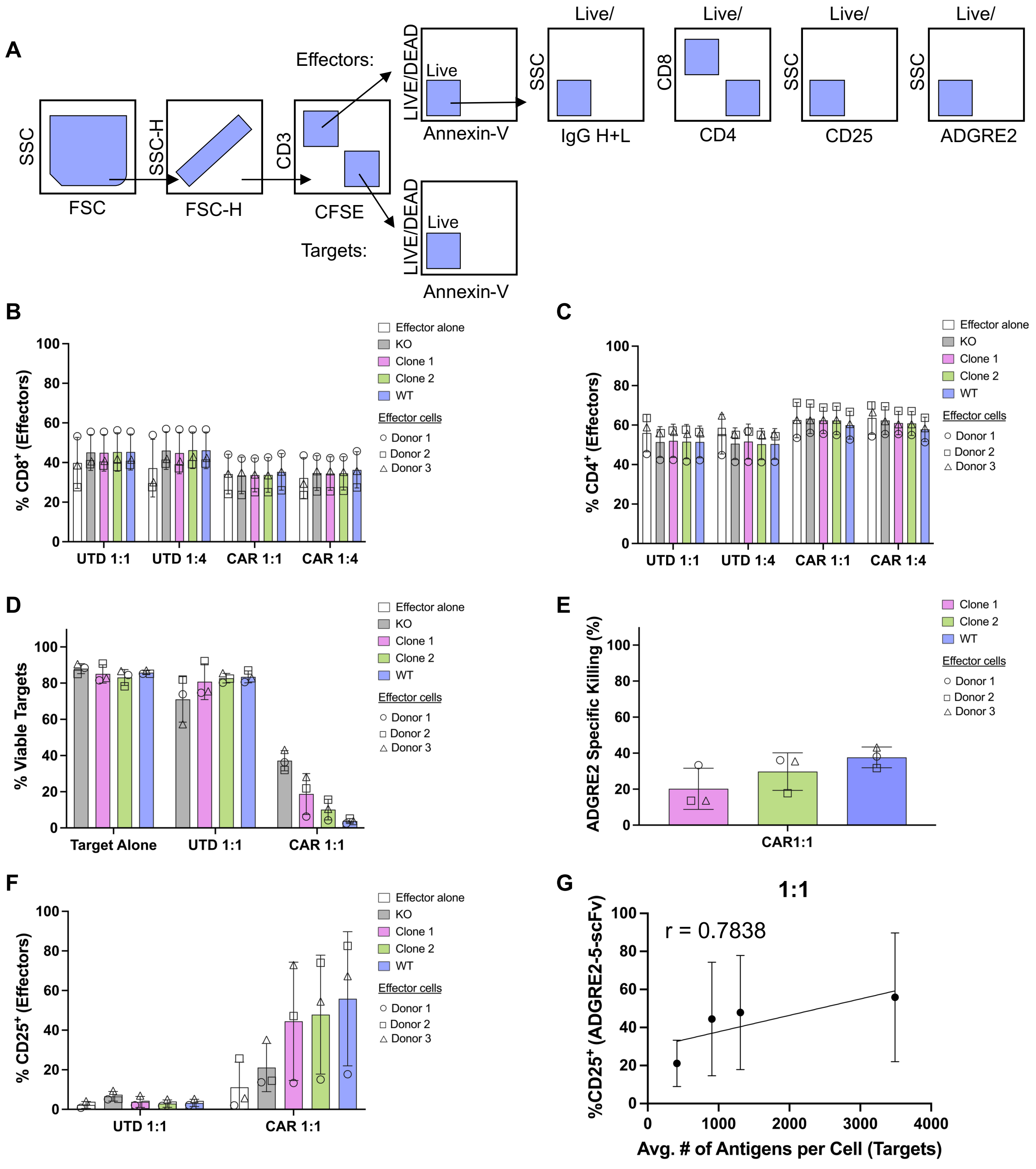
A
Effectors:
Targets:
SSC
FSC
SSC-H
FSC-H
CD3
CFSE
LIVE/DEAD
Live
Annexin-V
Live/
IgG H+L
CD8
CD4
CD25
ADGRE2
B
% CD8+ (Effectors)
UTD 1:1
UTD 1:4
CAR 1:1
CAR 1:4
Effector alone
KO
Clone 1
Clone 2
WT
Effector cells
Donor 1
Donor 2
Donor 3
C
% CD4+ (Effectors)
D
% Viable Targets
Target Alone
E
ADGRE2 Specific Killing (%)
CAR1:1
F
% CD25+ (Effectors)
G
1:1
r = 0.7838
%CD25+ (ADGRE2-5-scFv)
Avg. # of Antigens per Cell (Targets)

**Supplemental Figure 1. Phenotypic characterization of effector T cells and cytotoxicity with MOLM-13 target cells.**

**(A)** Flow cytometry gating strategy of co-culture readout. **(B-C)** Flow cytometric analysis of effector CD8 and CD4 surface expression after 48 h co-culture with MOLM-13 targets. Effectors cultured alone were included as controls. Data represented as mean ± SD from n=3 T cell donors. **(D)** *In vitro* cytotoxicity assessment of untransduced (UTD) or ADGRE2-5-scFv (CAR) T cells against MOLM-13 targets at a 1:1 effector-to-target (E:T) ratio. Viability was assessed after 48 h by flow cytometry (Annexin-V$^-$/LIVE-DEAD$^-$). Target-alone cultures served as controls. Data represented as mean ± SD from n=3 T cell donors. **(E)** ADGRE2-specific killing of MOLM-13 clones and WT at a 1:1 E:T ratio, calculated as $(Viability_{untreated} - Viability_{treated})/Viability_{untreated}\ x\ 100$ followed by subtracting killing observed in KO targets for background correction. Data represented as mean ± SD from n=3 T cell donors. **(F)** Effector activation measured by CD25 surface expression after 48 h co-culture with indicated MOLM-13 targets; flow cytometry analysis. Effector-alone condition included as a control. Data shown as mean ± SD from n=3 donors. **(G)** Correlation between ADGRE2-5-scFv T cell activation (CD25$^+$) and ADGRE2 surface intensity (antigens per cell, APC) on MOLM-13 targets at a 1:1 effector-to-target (E:T) ratio. Data shown as mean ± SD. Pearson correlation efficient (r) and linear regression analysis were generated using GraphPad Prism. Abbreviations: KO, knockout; SD, standard deviation.

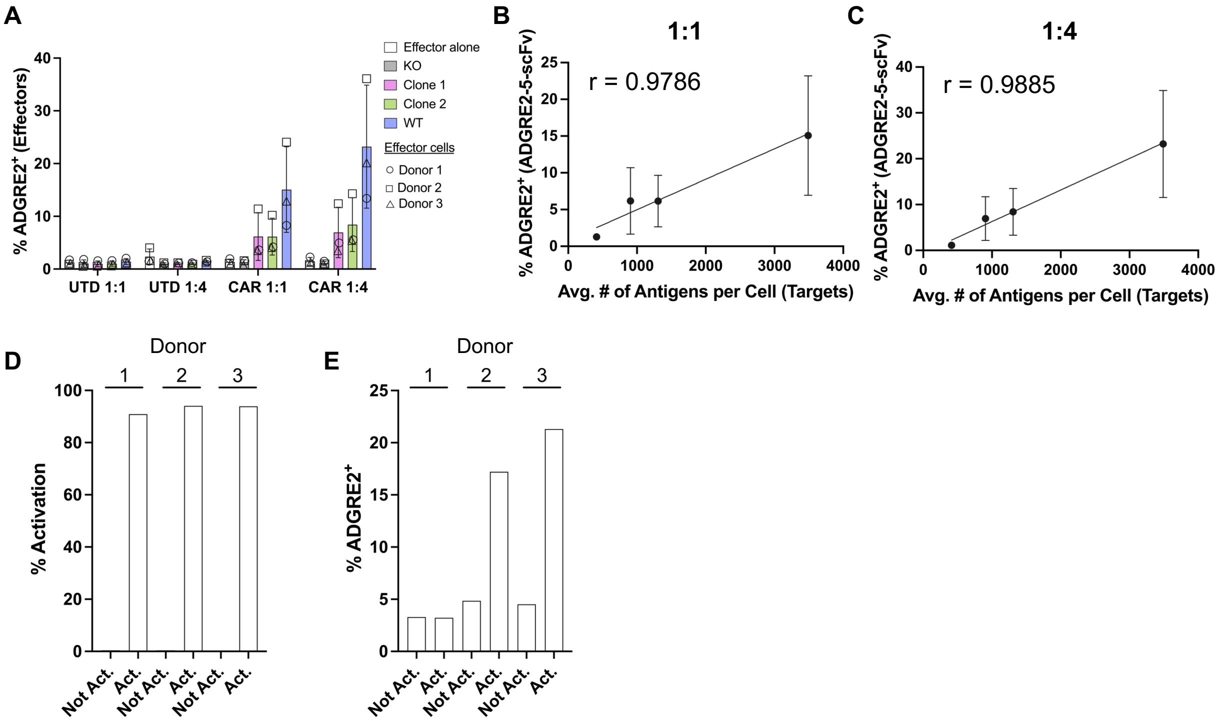

A
% ADGRE2+ (Effectors)
40
30
20
10
0
UTD 1:1
UTD 1:4
CAR 1:1
CAR 1:4
Effector alone
KO
Clone 1
Clone 2
WT
Effector cells
Donor 1
Donor 2
Donor 3
B
1:1
r = 0.9786
% ADGRE2+ (ADGRE2-5-scFv)
25
20
15
10
5
0
0
1000
2000
3000
4000
Avg. # of Antigens per Cell (Targets)
C
1:4
r = 0.9885
% ADGRE2+ (ADGRE2-5-scFv)
40
30
20
10
0
0
1000
2000
3000
4000
Avg. # of Antigens per Cell (Targets)
D
Donor
1
2
3
% Activation
100
80
60
40
20
0
Not Act.
Act.
Not Act.
Act.
Not Act.
Act.
E
Donor
1
2
3
% ADGRE2+
25
20
15
10
5
0
Not Act.
Act.
Not Act.
Act.
Not Act.
Act.

**Supplemental Figure 2. ADGRE2 expression on T cells following activation.**

**(A)** Flow cytometric analysis of ADGRE2 expression on effector T cells (UTD and ADGRE2-5-scFv CAR T (CAR) after 48 h co-culture at indicated effector-to-target (E:T) ratios. **(B-C)** Correlation between ADGRE2 surface expression on ADGRE2-5-scFv effector cells and ADGRE2 antigens per cell (APC) on MOLM-13 targets at 1:1 and 1:4 E:T ratios. Data represented as mean ± SD. Pearson correlation efficient (r) and linear regression analysis completed with GraphPad Prism. **(D)** T cell activation following CD3/CD28 bead stimulation. Percent $CD25^+$ and $CD69^+$ cells (within the $CD3^+$ population) were quantified by flow cytometry after 48 h in stimulated (Act.) or non-stimulated (Not Act.) conditions. Measured by flow cytometry; n=3 T cell donors. **(E)** ADGRE2 surface expression on non-activated (Not. Act.) or 48 h CD3/CD28 activated (Act.) pan-T cells, measured by flow cytometry. Data shown for n=3 T cell donors. Abbreviations: UTD, untransduced; SD, standard deviation.

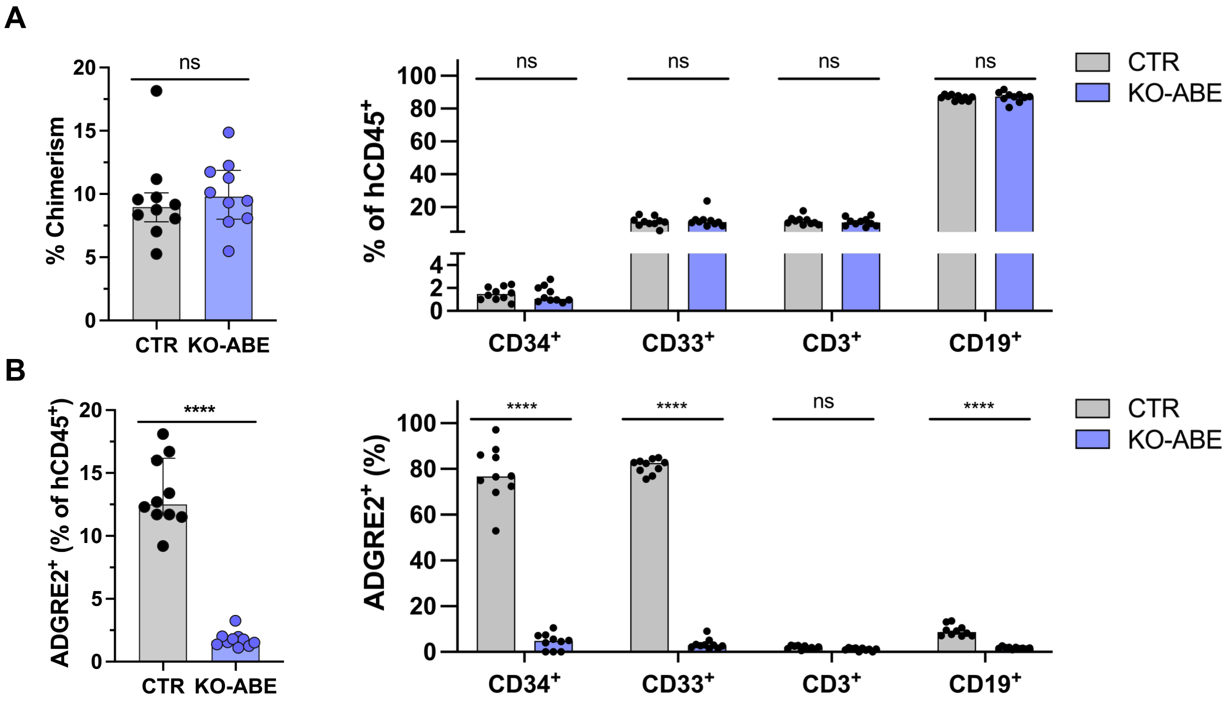
A
ns
% Chimerism
20
15
10
5
0
CTR
KO-ABE
% of hCD45+
100
80
60
40
20
4
2
0
ns
ns
ns
ns
CD34+
CD33+
CD3+
CD19+
CTR
KO-ABE
B
****
ADGRE2+ (% of hCD45+)
20
15
10
5
0
CTR
KO-ABE
ADGRE2+ (%)
100
80
60
40
20
0
****
****
ns
****
CD34+
CD33+
CD3+
CD19+
CTR
KO-ABE

**Supplemental Figure 3.** ***In vivo*** **pharmacology study peripheral blood analysis.** **(A)** Chimerism and multilineage reconstitution in peripheral blood of NSG mice, 16 weeks post-engraftment; measured by flow cytometry. Human chimerism calculated as $hCD45^+$ / ($hCD45^+$ + $mCD45^+$) x 100. Frequencies of $CD34^+$ (HSPC marker), $CD33^+$ (myeloid marker), and lymphoid (T cells, $CD3^+$; B-cell, $CD19^+$) populations were measured within $hCD45^+$ compartment. Data shown mean ± SD; n=10 mice per group. Each dot represents an individual mouse peripheral blood sample. Statistical analysis by two-way ANOVA between ADGRE2-KO and CTR mice; ns = not significant ($p > 0.05$). **(B)** ADGRE2 surface protein expression in total peripheral blood $hCD45^+$, $CD33^+$ myeloid, and lymphoid ($CD3^+$, $CD19^+$) populations in KO-ABE versus Control mice; measured by flow cytometry. Data represented as mean ± SD; n=10 mice per arm; each dot represents an individual mouse peripheral blood sample. Statistical analysis performed using two-way ANOVA; ns = not significant ($p > 0.05$), ****$p<0.0001$. Abbreviations: NSG, NOD-scid IL2Rg null; hCD45, human CD45; mCD45, murine CD45; SD, standard deviation; KO, knockout; CTR, control.
